# Allosteric inhibition of adenylyl cyclase type 5 by G-protein: a molecular dynamics study

**DOI:** 10.1101/2020.06.23.166710

**Authors:** Elisa Frezza, Tina-Méryl Amans, Juliette Martin

**Author notes:** (EF), (JM).

## Abstract

Adenylyl cyclases (ACs) have a crucial role in many signal transduction pathways, in particular in the intricate control of cyclic AMP (cAMP) generation from adenosine triphosphate (ATP). Using homology models developed from existing structural data and docking experiments, we have carried out all-atom, microsecond-scale molecular dynamics simulations on the AC5 isoform of adenylyl cyclase bound to the inhibitory G-protein subunit Gαi in the presence and in the absence of ATP. The results show that Gαi have significant effects on the structure and flexibility of adenylyl cyclase, as observed earlier for the binding of ATP and Gsα. New data on Gαi bound to the C1 domain of AC5 help to explain how Gαi inhibits enzyme activity and to get insight on its regulation. Simulations also suggest a crucial role of ATP in the regulation of stimulation and inhibition of AC5.

**Author summary:** The neurons that compose the human brain are able to respond to multiple inputs from other neurons. The chemical “integration” of these inputs then decides whether a given neuron passes on a signal or not. External chemical messages act on neurons via proteins in their membranes that trigger cascades of reactions within the cell. One key molecule in these signaling cascades is cyclic adenosine monophosphate (cAMP) that is chemically synthesized from adenosine triphosphate (ATP) by the enzyme adenylyl cyclase (AC). We are investigating the mechanisms that control how much cAMP is produced as a function of the signals received by the neuron. In particular, we have studied the inhibition effect of a key protein, termed Gαi, on AC, and we compare it with the stimulator effect of another key protein termed Gsα. Using microsecond molecular simulations, we have been able to show how binding Gαi to AC changes its structure and its dynamics so that its enzymatic activity is quenched and that ATP seems to have a crucial role in the regulation of stimulation and inhibition of AC5.

## Introduction

One of the most studied signal transduction pathways is the intricate control of cyclic AMP (cAMP) generation, a universal second messenger based on G-protein coupled receptors (GPCR) in eukaryotes [1]. cAMP has a role in a vast number of biological systems, including but not limited to hormone secretion [2], smooth muscle relaxation [3], olfaction [4], learning and memory [5–7].

The family of enzymes responsible for cAMP synthesis is the adenylyl cyclases (also commonly known as adenylate cyclases) which are highly regulated in order to tightly control cAMP levels [8]. Nine mammalian transmembrane ACs are recognized, with a cytoplasmic domain with catalytic properties (hereafter termed AC1-9) [8]. Each member of the family has specific regulatory properties and tissue distributions [9,10]; however they all convert adenosine triphosphate (ATP) into cAMP via a cyclization reaction.

Mammalian ACs share a similar topology of a variable N-terminus (NT) and two repeats of a membrane-spanning domain followed by a cytoplasmic domain [11,12]. The two cytoplasmic domains, called C1 and C2, contain a region of approximately 230 amino acid residues that are roughly 40% identical, called C1a and C2a: This implies a pseudosymmetry in ACs. Together the cytoplasmic domains form the catalytic moiety at the interface. The NT and C-terminal portion of the C1 and C2 domains, called C1b and C2b, are the most variable regions among the different isoforms and can differ among the species. The catalytic site of ACs is located at the C1/C2 interface and binds a molecule of ATP accompanied by two magnesium ions [13].

ACs’ function is regulated by several modulators, either stimulators or inhibitors of cAMP synthesis. These include the stimulatory G-protein subunit alpha (Gsα) which is released from its cognate receptor and binds to and activates the AC enzyme via the subunit interaction with the C2 domain [10,14–16] upon GPCR activation [14,16,17], the inhibitory G-protein subunits Gαi and Gβγ, calcium ions, calmodulin and a variety of kinases. AC isoforms integrate several signals and they differ from each other for their modulators and for the different tissues where they are more abundant [18–21]. Although all nine transmembrane ACs are expressed in the brain, specific ACs are particular abundant in specific brain regions, and AC5 is highly expressed in the striatum, and therefore involved in signal transduction networks that are crucial for synaptic plasticity in the two types of medium spiny neurons [22].

Structural information on AC cytoplasmic catalytic core [21] and on a complex containing both AC catalytic domains bound to an active conformation of the stimulating Gsα, with or without a bound ATP analog, is available [23]. However, the transmembrane regions contain six predicted membrane-spanning helices each and their function, aside from membrane localization, is unknown. Although the mechanism of stimulation of AC by Gsα is relatively well understood, the mechanism of inhibition of AC activity is still debated: some mutational studies suggest that Gαi binds in an opposite binding site [24], but there are other hypotheses, like the possibility of simultaneously binding of Gαi and Gsα or a competition between the two G proteins. However, there are no data on the enzyme bound to ATP (or an ATP analog) in the absence of activating Gsα, on the enzyme in complex with Gαi in the presence and absence of ATP and on the possible trimeric form Gαi+AC+Gsα in the presence or absence of ATP. Hence, it is difficult to understand how Gα subunits activate/inhibit adenylyl cyclase and what is the role of ATP.

To gain insight into the functional mechanism of AC, some studies at the molecular level have been conducted using all-atom molecular dynamics (MD) simulations. In our previous work, we studied the stimulation mechanism of AC5, by performing MD simulations of AC5 alone, AC5+ATP, AC5+Gsα and AC5+ATP+Gsα [25]. We chose the mouse AC5 isoform among the other isoforms, since this isoform notably plays a key role in a variety of neuronal GPCRs-based signal cascades [19,26,27]. We extensively characterized the flexibility of the four states, the protein-protein interfaces, the ATP mobility, the Gsα binding site and the Gαi putative binding site on C1 and the effect of the ATP and Gsα on these properties. Our study showed that both ATP and Gsα binding have significant effects on the structure and flexibility of adenylyl cyclase. The comparison between the simulations of AC5+ATP and AC5+ATP+Gsα helped to explain how Gsα binding enhances enzyme activity and could therefore aid product release. Our simulations also suggested that ATP binding could influence the binding of the inhibitory G-protein subunit Gαi, if the potential binding site within domain C1 were to be involved.

At the same time, another study by Van Keulen and co-workers has been published where they investigated the mechanism of inhibition of AC5 by N-terminal myristoylated Gαi. In their studies, they considered apo AC5 (i.e. without ATP) and they concluded that the myristoylation seems to play a crucial role for the inhibition of AC5. By binding Gαi to a postulated C1 binding site, they found structural modifications that would disfavor both ATP and Gsα [28]. Recently, they have also characterized the complex Gαi+AC5+Gsα using N-terminal myristoylated Gαi [29,30] by comparing the different simulations in order to understand the impact of the binding of both Gα proteins. This comparison suggests that association of both Gαi and Gsα subunits results in an AC5 conformation similar to that sampled by the Gαi+AC5 complex, indicating that the ternary complex mainly samples an inactive conformation.

Despite these recent studies, what impact Gαi would have if ATP were already bound in its AC5 pocket and also whether Gsα and Gαi could nevertheless bind simultaneously to AC5 in the presence of ATP is yet to be clarified. In the present study, we used the same approach applied to investigate the stimulation mechanism in our previous work [25]. We have used all-atom molecular dynamics simulations to study the impact of ATP and Gαi on the structure and flexibility of AC5. As in our previous study, we considered only the cytoplasmic domains of AC5, since they are capable of reproducing many of the regulatory properties of the wild type enzyme and therefore can be used as working models to investigate the regulation mechanisms of AC [31,32]. Since no structural data are available for the complex AC5+Gαi, we computed docking experiments using representative structures for Gαi and AC5+ATP obtained from our MD simulations and we considered two distinctive poses. The all-atom microsecond-scale simulations of AC5 in complex with Gαi with or without ATP studied here (see Figure 1) were compared with our previous simulations of AC5, AC5+ATP, AC5+Gsα and AC5+ATP+Gsα in order to help to explain how binding changes the properties of AC5 and notably to understand the inhibition effect of Gαi.

**Figure 1.**
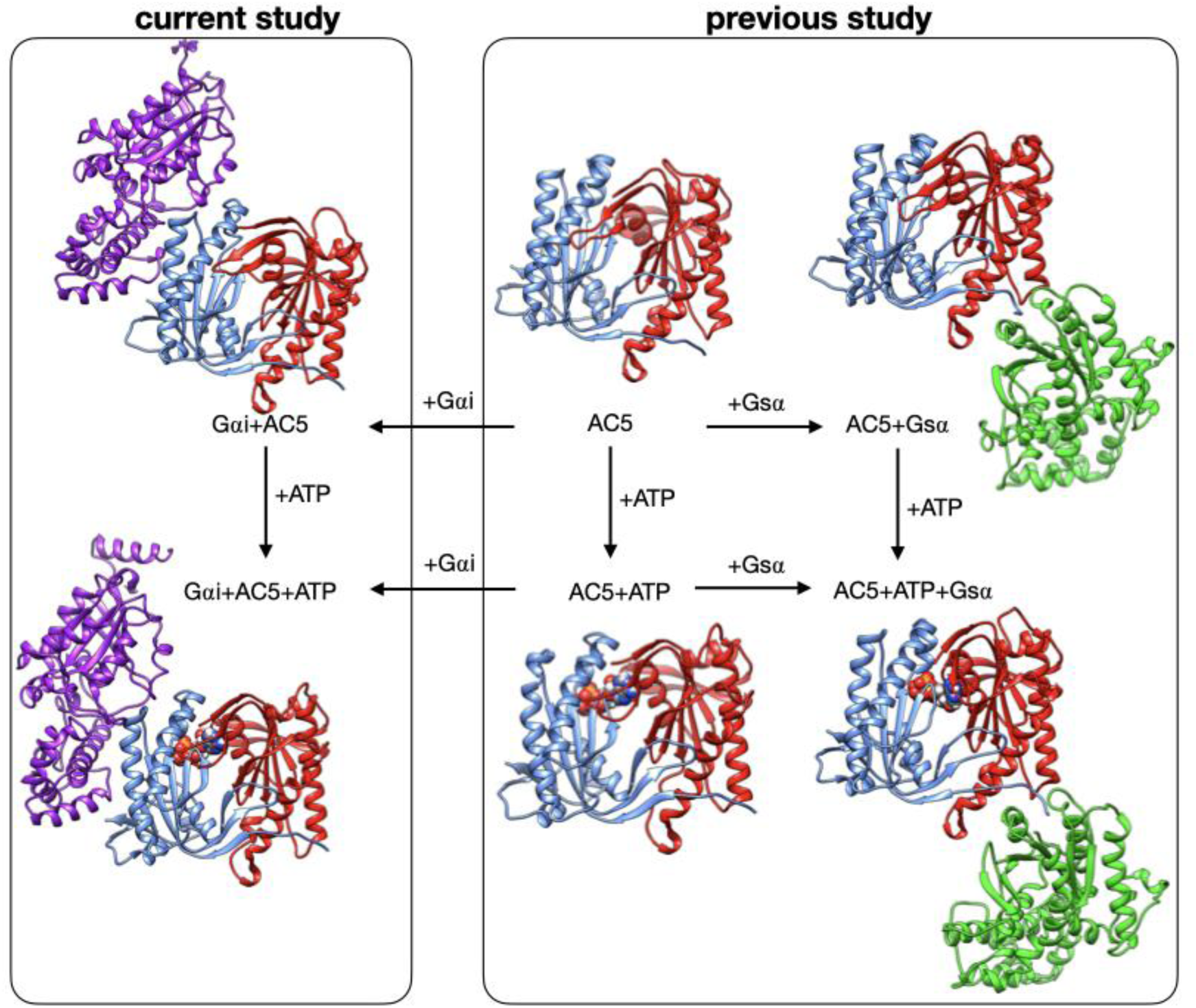
Structure of the cytoplasmic segment of the AC5 isoform of adenylyl cyclase and of its complexes with ATP and the regulating G-proteins Gsα and Gαi viewed from the side closest to the cell membrane. Proteins are shown as backbone ribbons. The C1 and C2 subunits of AC5 are colored blue and red respectively, Gsα is colored green and Gαi is colored purple. ATP is shown in a CPK representation with standard chemical coloring. In each case, the structures are averages taken from the molecular dynamics simulations. For the AC5 in complex with Gαi with and without ATP, we chose one of the docking poses we used in this work.

## Results

### Overview of simulations

In the absence of any structural information on the catalytic domains of the enzyme with the inhibiting G-protein subunit Gαi, we use a combination of homology modelling, molecular dynamics and protein-protein docking to get insight on the inhibition mechanism at molecular level and the impact of the ligand or protein on the conformation and dynamics of AC5.

We have studied the behavior of two molecular species (see Figure 1): AC5 bound to the inhibiting G-protein subunit Gαi (Gαi+AC5) and AC5 bound to both ATP and Gαi (Gαi+AC5+ATP). For both complexes, we considered two different relative conformations: one called Gαi_sym+AC5 where the Gαi protein has an orientation symmetrical to the Gsα protein in the AC5+Gsα complex, presented in Figure 2A, and one called Gαi_tilted+AC5, where the Gαi protein is tilted, presented in Figure 2B. For each of these species, we generated 1.5 µs MD trajectories in an aqueous environment with a physiological salt concentration (0.15 M KCl) using the GROMACS 5 package [33–36] with the Amber 99SB-ILDN force field for proteins [37,38]. The first 400 ns of each trajectory were treated as equilibration of the system and analysis was carried out only on the remaining 1.1 µs. We analyzed all-atom MD simulations using average structures, time-averaged properties, angles and distances between helices (see Figure 3), specific geometrical measurements to characterize protein-protein and protein-ligand interface and residue-by-residue conformational and dynamic properties. In order to understand the effect of Gαi and ATP on AC5, we used the MD simulations for isolated AC5, AC5 with ATP and two Mg2+ ions in its active site (AC5+ATP), AC5 bound to the activating G-protein subunit Gsα (AC5+Gsα+GTP) and AC5 bound to both ATP and Gsα (AC5+ATP+Gsα+GTP) obtained in our previous work [25,39]. Data are shown in Table S1-S3 and in Figure S3-S12.

**Figure 2.**
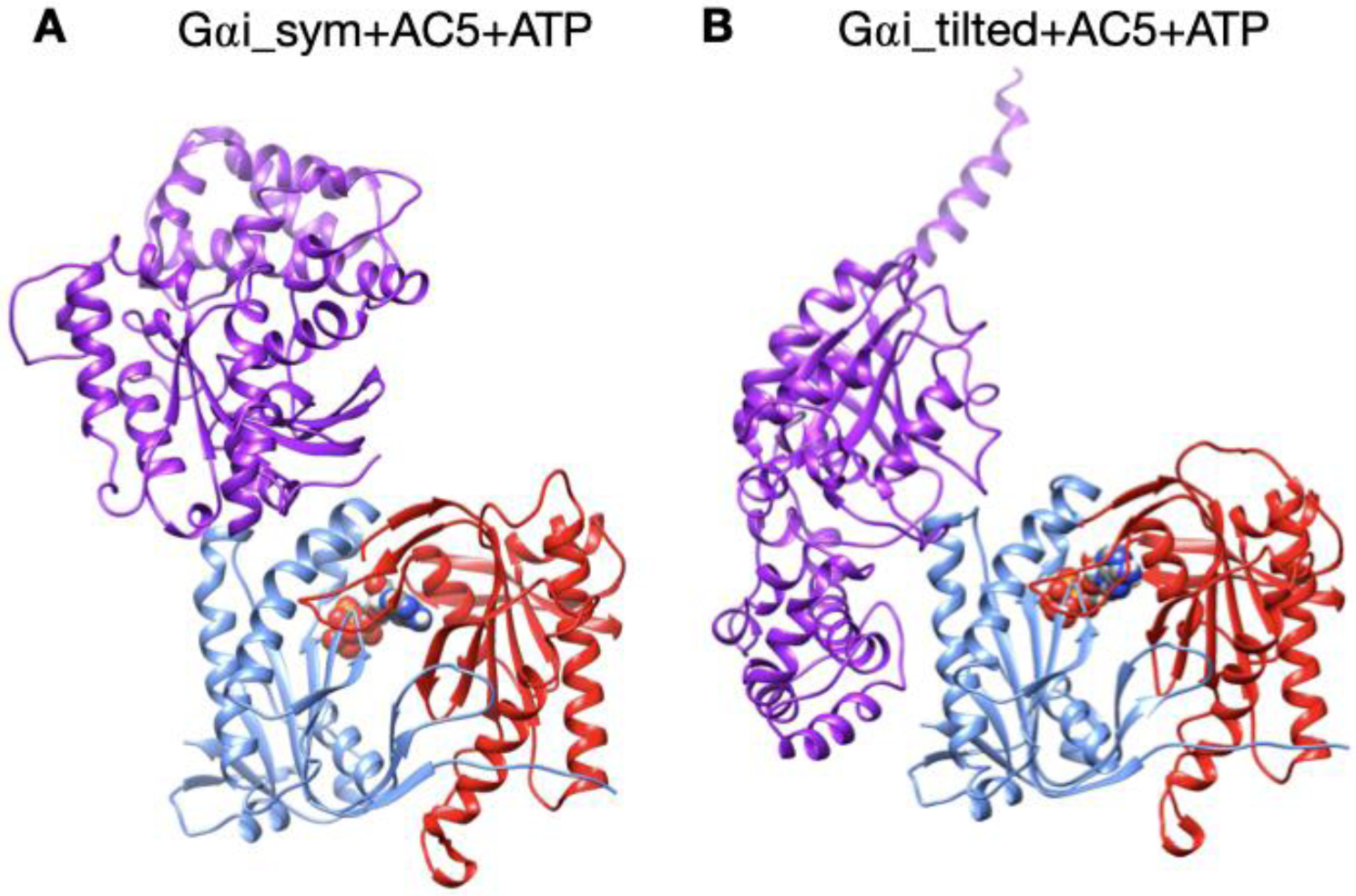
The two different configurations of the Gαi+AC5+ATP complex simulated in this study. A: Gαi_sym+AC5+ATP: Gαi has an orientation symmetrical to the Gsα protein in the AC5+Gsα complex. B: Gαi_titlted+AC5+ATP: Gαi protein is tilted with respect to AC5.

**Figure 3.**
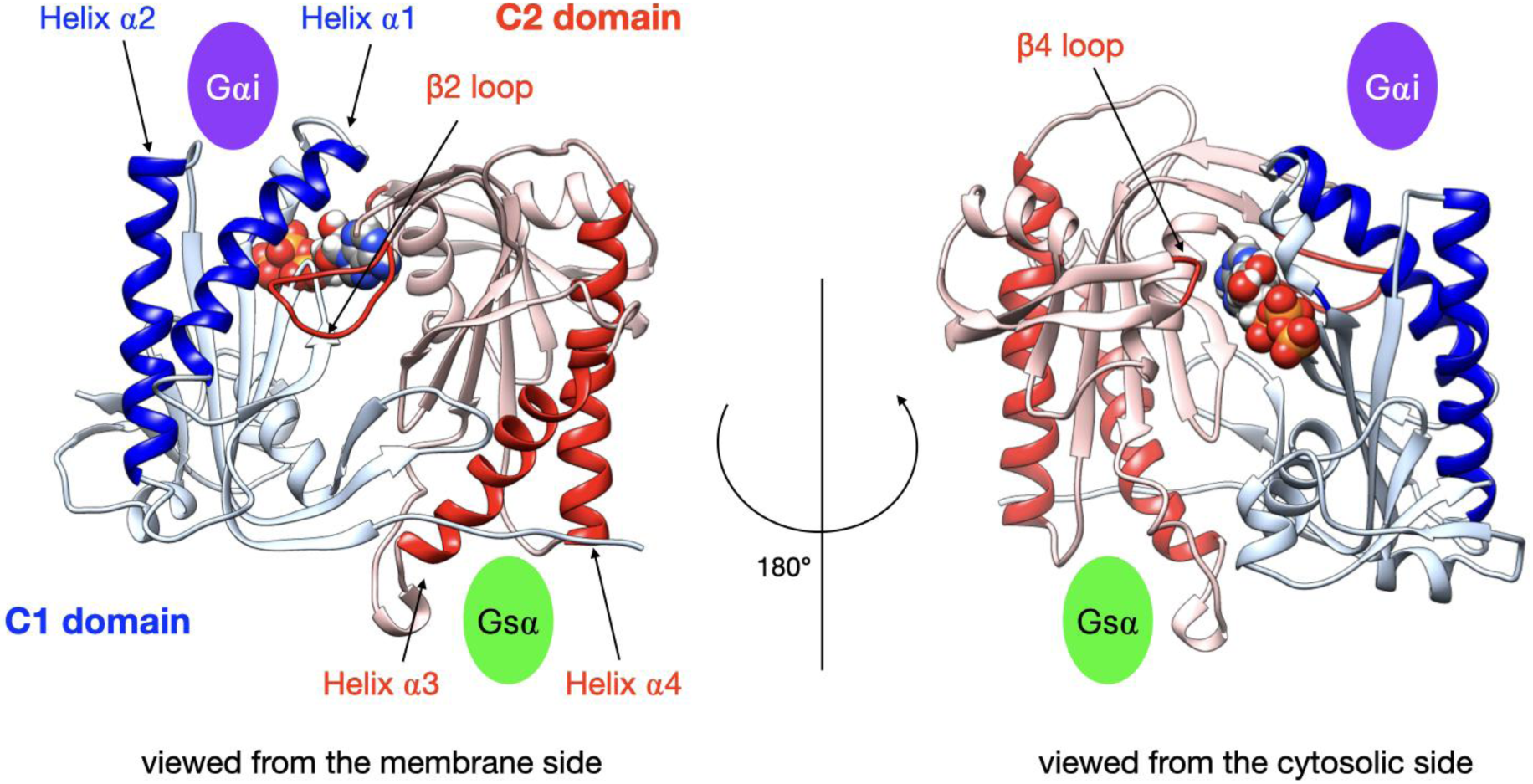
Illustration of the key regions of AC5 catalytic domain structure, with bound ATP. The C1 domain is colored blue and the C2 domain in red, with relevant parts in darker color: the helices of C2 involved in binding of the stimulatory protein Gsα, the helices of C1 involved in binding of the inhibitory protein Gαi, the β2 loop of C2 (left side) and the β4 loop of C2 (right side) which bears the catalytic Lysine residue. The green oval indicates the binding site of Gsα and the purple oval indicates the binding site of Gαi.

### Stability of Gαi+AC5 complexes in the presence and in the absence of ATP

The two types of complexes behave differently in the presence of ATP. In the Gαi_sym+AC5+ATP simulation, the Gαi protein reallocates significantly with respect to AC5 toward the C2 domain, ending in a configuration where it is in contact with the C2 domain, see Figure 4A. The peculiarity of this system is also apparent in the rest of the study and will be commented later. On the contrary, in the Gαi_tilted+AC5+ATP system, the Gαi protein fluctuates around its initial position, without significant reallocation, indicating that this complex is very stable, see Figure 4B. We quantified the fraction of interface contacts between Gαi and AC5 that are conserved throughout the simulation time, and the total number of interface contacts as a proxy of the interface size. For Gαi_sym+AC5+ATP, the fraction of conserved contacts at the Gαi/AC5 interface drops rapidly below 25%, while the total number of contacts increases significantly (see Figure S1). In Gαi_tilted+AC5+ATP, 50 to 75% of the initial contacts are conserved during the simulation time, and the total number of contacts also tends to increase, although less dramatically (Figure S1). In the absence of ATP, both systems maintain between 50 and 75% of their initial contacts, with more moderate reallocation and variation in terms of contact number (see Figure S1 and S2).

**Figure 4.**
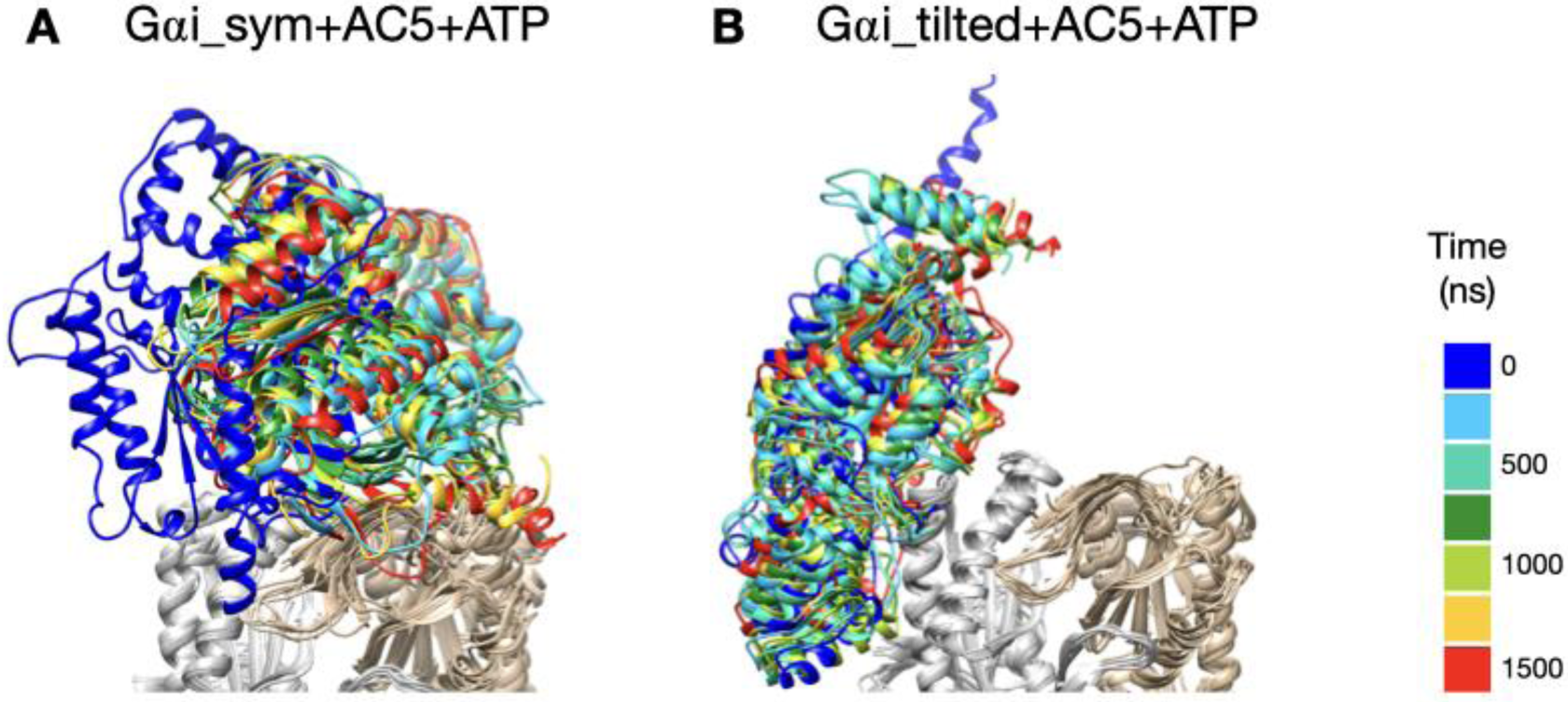
Snapshots of the Gαi+AC5+ATP complexes observed during the simulations, viewed from the membrane side. Gαi structures extracted every 250 ns are colored on a rainbow scale from blue to red. The C1 domain of AC5 is colored in grey and the C2 domain in beige.

### Impact of Gαi on AC5+ATP

We begin by considering the global impact of Gαi on AC5+ATP by computing the RMSD on backbone atoms separately on each AC5 domain. RMSD calculations with respect to the average MD structure of each AC5 domain show that Gαi binding has a significant effect on both the structure and the dynamics of the enzyme (see Table S1, and Figure S3 where in order to allow comparison with the results obtained in our previous study [25] values for AC5, AC5+ATP, AC5+Gsα, AC5+ATP+Gsα were also included). On the one hand, the domain C1 is slightly rigidified by the binding of Gαi. On the other hand, the C2 domain visits several conformational substates involving the ATP binding pocket (β2 loop and β4 loops) in Gαi_sym+AC5 complex (see Figure 5). These two substates also lead to two different substates for ATP (see Figure S4) which is more mobile, increasing the average RMSD from 0.6 Å for AC5+ATP to 0.9 Å for Gαi_sym+AC5+ATP (see Table S3 and Figure S5). In the case of Gαi_tilted+AC5+ATP, the C2 domain visits a specific substate close to the one sampled in AC5+ATP where the mobility of ATP is unchanged due to the presence of Gαi.

**Figure 5.**
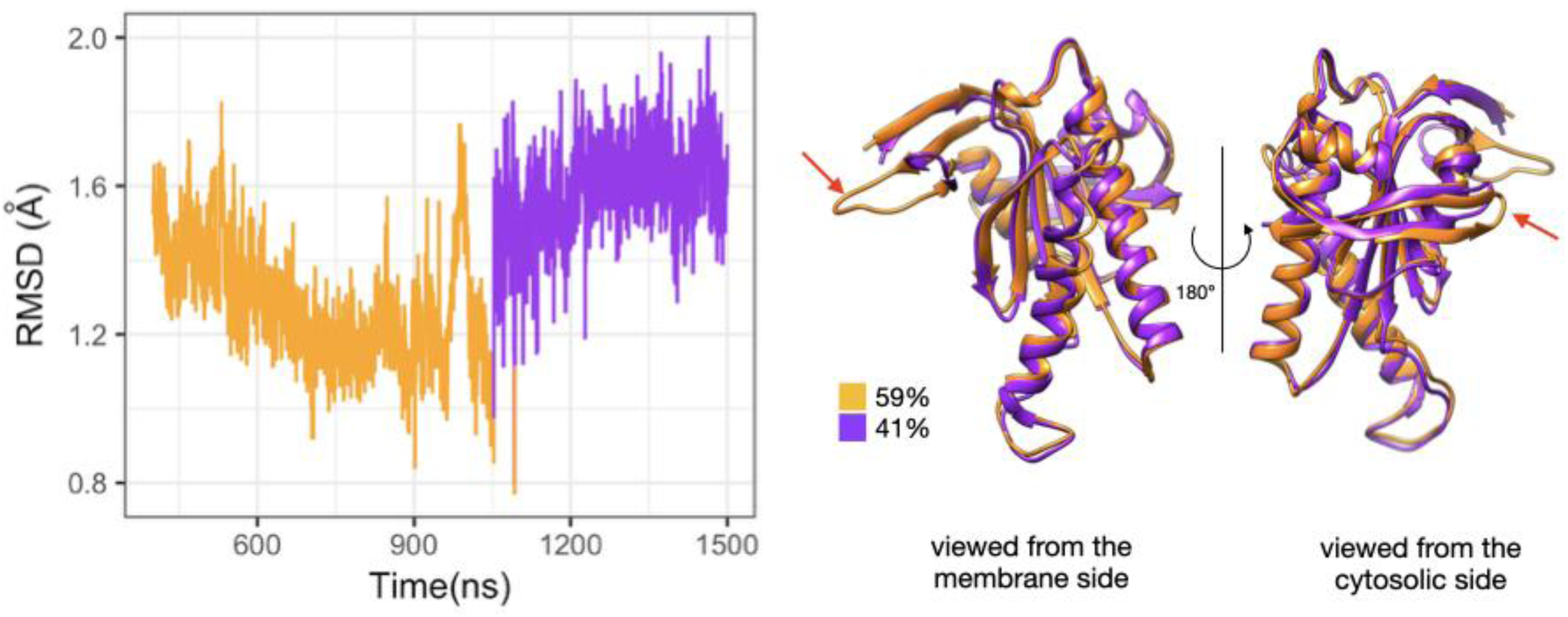
Sub-states of domain C2 observed during the simulation of Gαi_sym+AC5+ATP complex. Left side: RMSD time series for the C2 domain, colored according to cluster membership. Right: structures closest to the center of each cluster, and relative size of each cluster as percentages. Prominent structural changes are indicated by red arrows.

The presence of several substates upon binding of Gαi is in contrast with the stabilization of a specific substate upon binding of ATP and/or Gsα. Indeed, in our previous work, we observed that the C2 domain can investigate several substates when AC5 is isolated, and the presence of either ATP alone or ATP and Gsα stabilizes two distinct substates. In the former case, a single substate for the β2 loop is selected (the longest-lived substate in isolated AC5) in a close conformation. In the latter, an opening of loop β2 away from the active site is observed. The selection of a specific substate is correlated to the mobility of ATP and its reactivity [25].

Despite the decrease in flexibility of the C2 domain, also observed when Gαs is bound to AC5+ATP, in the presence of Gαi, ATP is still rather mobile (see Table S3) and for Gαi_sym+AC5+ATP an increase in mobility is observed: this impact is opposite to the one observed in AC5+ATP+Gsα, where a higher stability of ATP is observed (average RMSD equal to 0.3 Å). In both simulations in the presence of ATP, the interactions between the terminal phosphate group of ATP and Lys 1065 (belonging to loop β2) and the interactions between the penultimate phosphate and Arg 1029, a key functional residue, are absent (see Figure S6): the arginine side chain is separated from its target oxygen atom by roughly 9 Å. Moreover, it is known that ATP has stronger interactions with the C1 domain via its associated two Mg^2+^ ions, notably with residues Asp 396 and Asp 440. For Gαi_tilted+AC5+ATP, these interactions are stable and are not affected by the presence of Gαi. On the contrary, for Gαi_sym+AC5+ATP, these interactions are absent justifying the increase of ATP mobility (see Table S3).

Gαi binding also turns out to have more global effects on AC5+ATP. First, the angles and the distance between the pairs of α-helices in both AC5 domains are modified. In Gαi_sym+AC5+ATP, the angle between the helices α1 and α2 in domain C1 is significantly reduced (by 9°, see Table S1) and the angle the helices α3 and α4 in domain C2 is slightly increased (by 4°). In Gαi_titled+AC5+ATP, an opposite effect is observed: the angle between the helices α1 and α2 in domain C1 is slightly affected (increased by 1°, see Table S1 and Figure S7) and the angle between the helices α3 and α4 in domain C2 is slightly decreased (by 3°). In addition, the distance between the C2 helices in Gαi_titled+AC5+ATP complexes is maintained around 13 Å, whereas our earlier results indicate that it is around 16 Å in the AC5+ATP+Gsα complex (see Table S1 and Figure S8).In both simulations, the C1/C2 interface remains mostly as tight as in isolated AC5+ATP (gap index from 2.8 Å for AC5+ATP to 2.9 Å once Gαi is bound, Table S1 and Figure S9) involving a movement of helix α3 (see Figure 6). In terms of flexibility, Gαi binding mainly flexibilizes the binding site region of ATP in AC5, although it also decreases the flexibility of the C-terminals of helices α1 and α3 (see Figure 7A).

**Figure 6:**
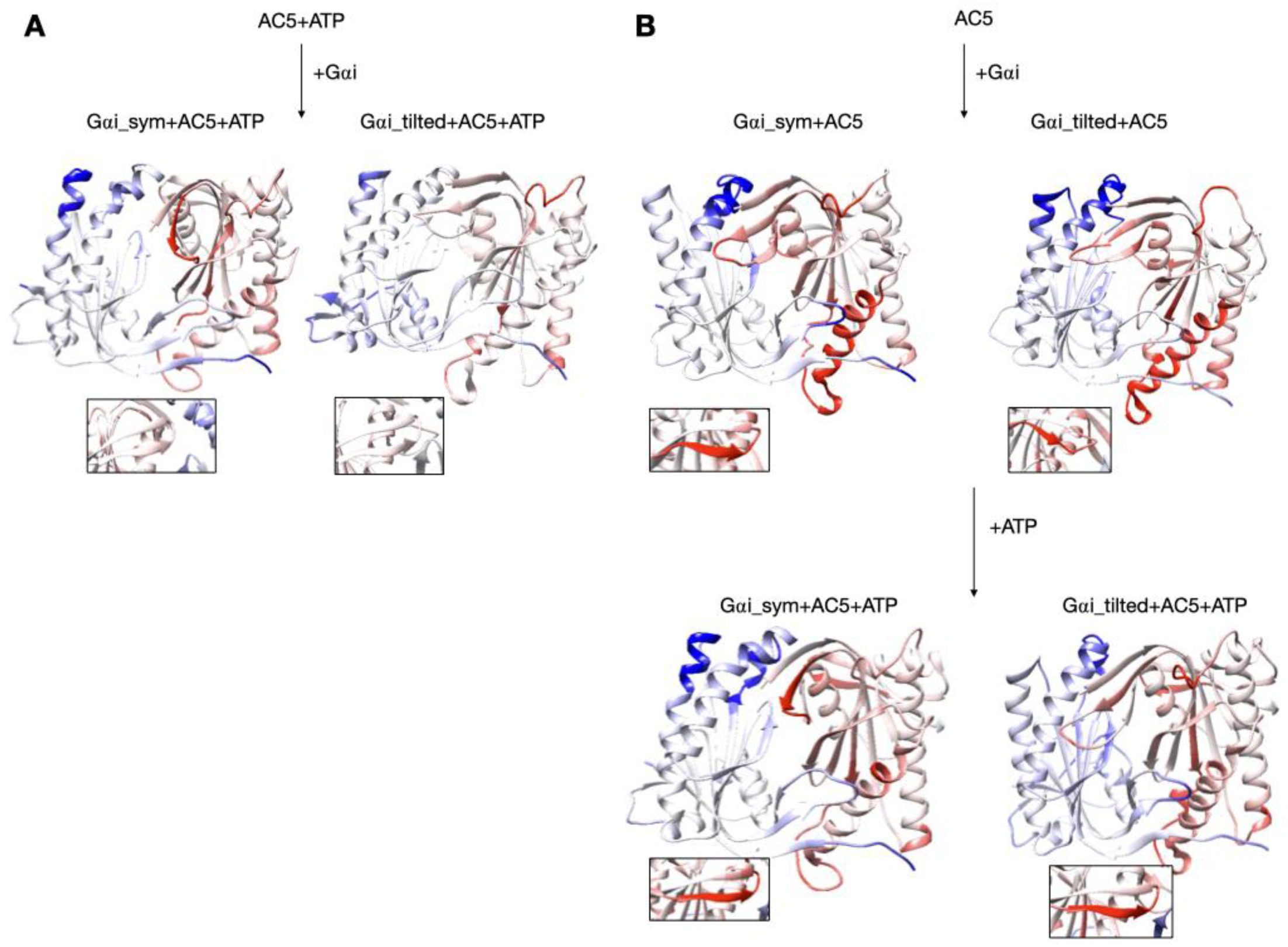
Changes in conformation induced by Gαi protein. A: scenario where ATP is already bound to AC5 when Gαi interacts, B: scenario where ATP is not yet bound to AC5 when Gαi interacts. More intense colors (blue for domain C1 and red for domain C2) correspond to larger movements compared to the preceding structure (i.e. AC5+ATP for A and AC5 and AC5+Gαi for B) on a scale of 0 to 4 Å. The insets display the β4 loop.

**Figure 7:**
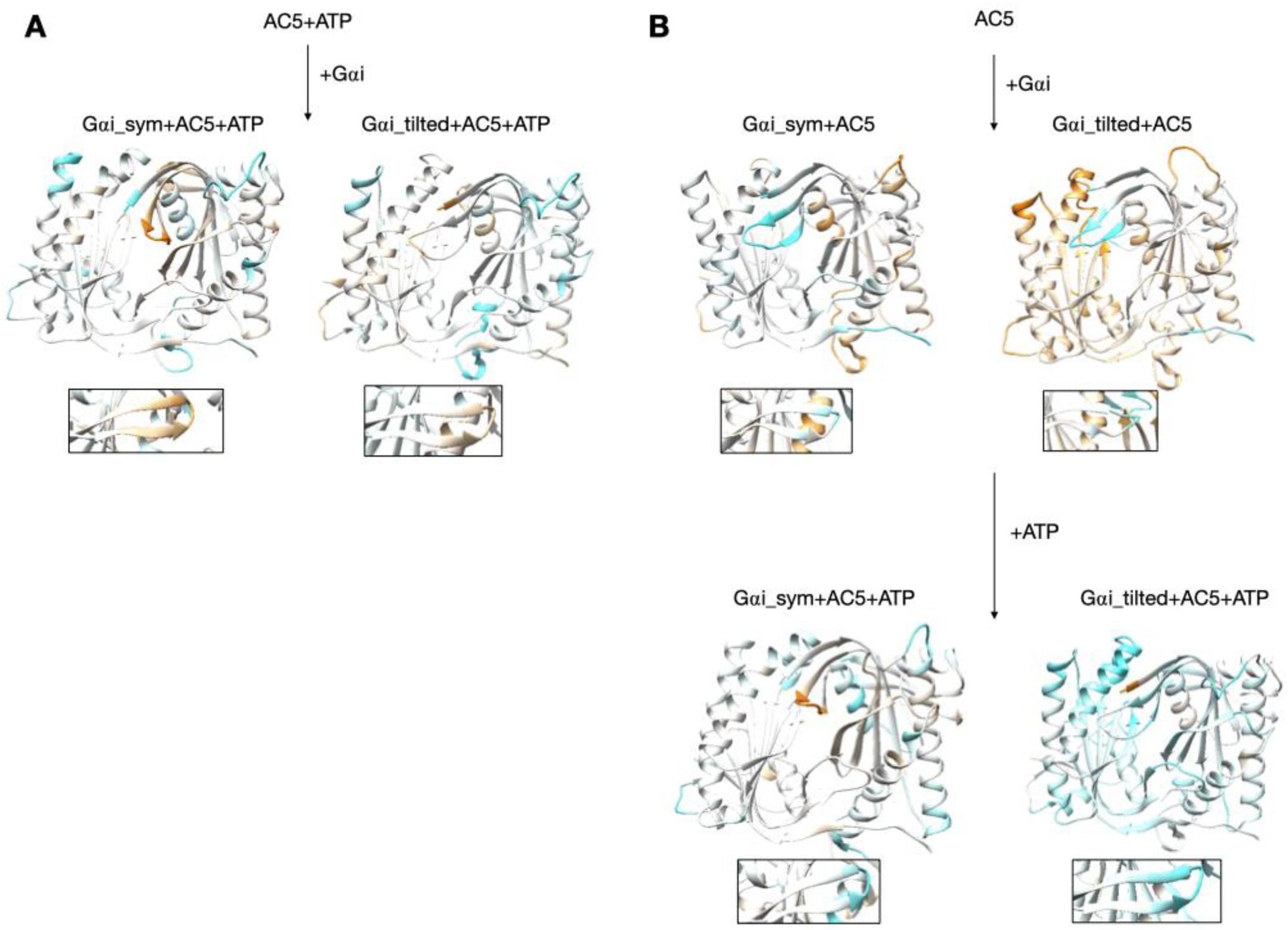
Changes in flexibility induced by G proteins. A: scenario where ATP is already bound to AC5 when Gαi interacts, B: scenario where ATP is not yet bound to AC5 when Gαi interacts. More intense colors (orange for increased flexibility and cyan for decreased flexibility) correspond to differences with respect to the preceding structure on a scale of −1.2 to +1.2 Å. The insets display the β4 loop.

### Impact of Gαi on *apo* AC5 and further impact on ATP on AC5+Gαi

Although it seems probable that ATP is already bound to AC5 based on our previous study [25], we also consider the scenario where ATP is not already present when Gαi binds on AC5. We begin by considering the global impact of Gαi on AC5. Gαi_sym and Gαi_tilted have different effects on the C1 domain of AC5: Gαi_sym slightly rigidifies it as attested by the RMSD calculation (Table S1 and Figure S3), whereas Gαi_tilted flexibilizes it, and this flexibility concerns the binding helices α1 and α2 (Figure 7B). For the C2 domain, in both Gαi_sym+AC5 and Gαi_tilted+AC5 complexes, the β2 loop and the β4 loop are rigidified upon addition of Gαi (see Figure 7B). Their conformations slightly differ in both complexes: the β2 loop is more closed with Gαi_tilted than with Gαi_sym whereas the β4 loop, at the back of the structure, is half open with Gαi_tilted compared to with Gαi_sym (see Figure S10). In Gαi_sym+AC5, the C2 domain visits several conformational substates involving the helices α3 and α4 (see Figure S11).

The conformation of the binding helices in both AC5 domains is significantly altered by Gαi. Gαi notably displaces helix α3 (Figure 6B). In both Gαi_sym+AC5 and Gαi_tilted+AC5, the angle between the helices α1 and α2 in domain C1 is significantly increased (by 24° and 19°; see Table S1 and Figure S7). On the contrary, the angle between the helices α3 and α4 in domain C2 is significantly decreased (by 7° and 12°), and the helices are also closer to each other (Table S1 and Figure S8). The C1/C2 interface remains as tight as in isolated AC5, in contrast with what we observed previously with the binding of Gsα, which resulted in a looser C1/C2 interface (gap index equal to 3.8 Å, see Table S2 and Figure S9).

In the scenario where ATP is not already bound to AC5 when Gαi interacts, we can also analyze the effect of ATP addition on the pre-formed Gαi+AC5 complex. As shown in Figure 7B, the addition of ATP notably rigidifies AC5. The interface at the Gαi/AC5 interface is quite loose in the Gαi_tilted+AC5 complex (gap index equal to 4.6 Å, see Table S2) and the addition of ATP tends to tighten this interface (gap index equal to 4.2 Å). On the contrary, in the Gαi_sym+AC5 complex, the initial Gαi/AC5 interface is made more loose by the addition of ATP (gap index from 5.4 Å in Gαi+AC5 to 3.2 Å when ATP is bound), probably due to the change of interface as observed by the losing of 75% of the native contacts by adding ATP (see Figure S1). By comparison, in the AC5+Gsα complex, the gap index decreased from 3.2 Å to 2.7 Å upon ATP addition (Table S2). Despite the variation of the values of gap index for the Gαi/AC5 interface upon binding of ATP, all the values are typical of obligate protein-protein interfaces [40].

## Discussion

As already observed in our previous work [25], microsecond-scale simulations are necessary to investigate the allosteric coupling existing within AC5 and the effect of the binding of Gαi. As van Keulen and Rothlisberger [28], we studied the scenario where Gαi binds to AC5 in the absence of ATP, but we could not exclude the possibility that ATP is bound on AC5 when Gαi binds, based on our previous work [25]. For the lack of structural information on the complex Gαi and AC5, we considered two different docking poses in our study: Gαi_sym and Gαi_tilted. In the former, the Gαi protein is bound to AC5 in a symmetrical fashion compared to what is known for the AC5+ATP+Gsα complexes (Figure 2A). In the latter, the Gαi protein is rotated and tilted onto the C1 domain (Figure 2B). Both complexes stay bound during the simulations, but a greater stability is obtained for the Gαi_tilted configuration, suggesting more biological relevance. In the case where Gαi binds on AC5+ATP in a symmetrical fashion, an allosteric effect is observed: a closure on the Gαi site is coupled with an opening on the Gsα site and an opening of the β2 loop of C2, as observed for Gsα. Despite that, this conformation seems to be less likely because the complementarity is very low and the interface is unstable, as already mentioned above. On the contrary, the Gαi_tilted configuration is also very similar to the one reported by van Keulen and Rothlisberger in their recent work where they studied the complex between myristoylated Gαi and AC5 in the absence of ATP [28], although the starting conformation of AC5 and Gαi are quite different. The binding of Gαi slightly rigidifies the C2 domain in all simulations and an opening of Gαi binding site is coupled with a closure of the Gsα binding site in particular in the Gαi_tilted+AC5 complexes with and without ATP, as observed in van Keulen and Roethlisberger’s simulation [28]. As in the case of AC5+Gsα complex, ATP stabilizes the Gαi/AC5 interface when Gαi is tilted, whereas the latter is less complementary. All these changes involve coupling through AC5 over distances of tens of angstroms.

Turning now to the enzymatic function of AC5, it is known that specific hydrogen bonds between AC5 and ATP play an important role in the production of cAMP from ATP as already shown in hybrid QM/MM free energy calculations, notably the hydrogen bond between the highly conserved Arg 1029 and the primary phosphate group of ATP [41]. The present simulations show that this interaction is not formed upon binding of Gαi and other ones are lost (for example between Lys 1065 and ATP, see Figure S11), when on the contrary it is formed upon binding of Gsα, see Figure 8.

**Figure 8:**
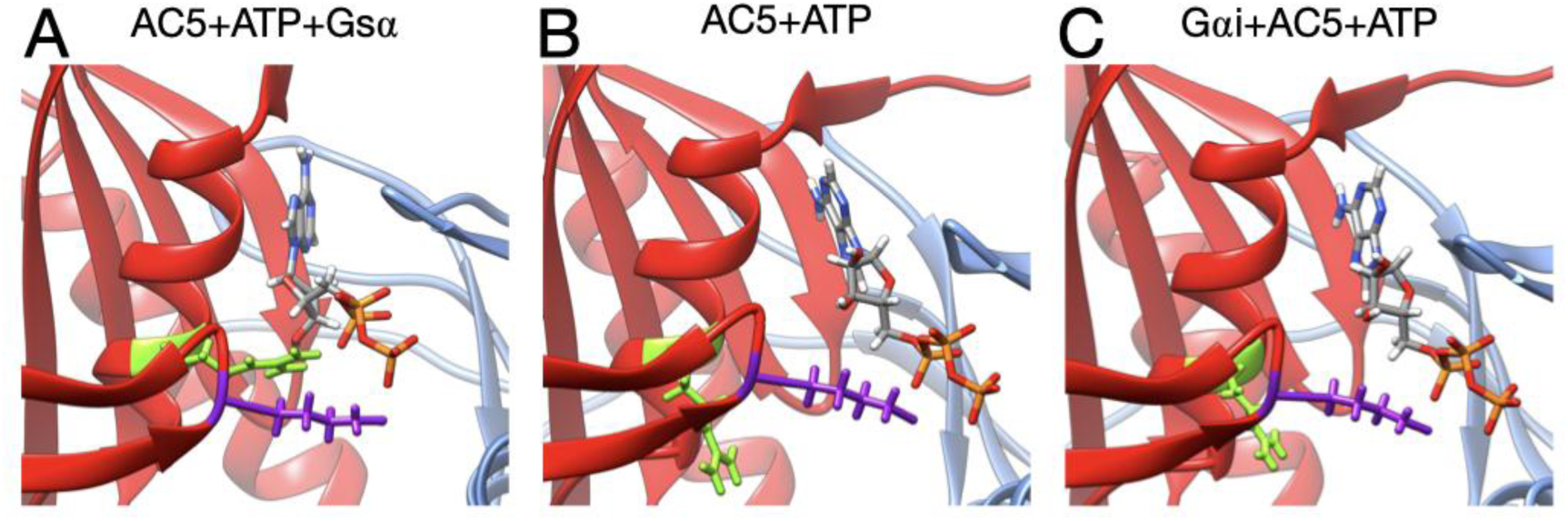
Interactions between ATP and key residues in different complexes. A: active AC5+ATP+Gsα, B: inactive AC5+ATP complex, C: inactive Gαi_tilted+AC5+ATP complex. L of C2 are shown as sticks and colored in purple (Lys 1065) and green (Arg 1029). For clarity, the region 394-428 of C1 has been omitted from the representation.

The binding of Gαi inhibits AC5 by increasing the flexibility of the active site allowing a high mobility of ATP without changing the complementarity of the C1/C2 interface. The further inactivation of AC5+ATP by Gαi does not allow to exclude the possibility that ATP is bound to AC5 during the inhibition process and it may have a role in the tight regulation of the enzyme.

Based on our previous study [25] and the current one, we can speculate on the regulation mechanism of AC5 by considering only the Gαi_tilted configuration. In the absence of ATP, based on our simulations, only Gsα could interact with AC5, due to the close conformation of the binding site on the C1 domain. However, we cannot exclude the existence of Gαi+AC5 based on our study. The binding of ATP induces an opening of the angle between the pair of α-helices of domain C1 (α_C1_) and a closure of the pair of α-helices of domain C2 (α_C2_), which have a similar value close to the one observed upon binding of Gsα. If Gαi binds first AC5+ATP, the closure of the angle α_C2_ does not allow the binding of Gsα and the ATP pocket remains close. When Gαi dissociates from AC5+ATP, AC5 undergoes a conformational change that allows the binding of Gsα. On the contrary, if Gsα binds first AC5, the enzyme is active thanks to the stabilization of ATP in its pocket and the formation of specific hydrogen bonds and cycles between AC5+ATP+Gs (favourable to Gi binding) and AC5+Gs (unfavourable to Gi binding). If Gαs dissociates, after cAMP release, AC5 is an apo conformation and there is no need for further inhibition. If Gαs dissociates from AC5+ATP, then the conformation of AC5 becomes accessible to Gαi for the inhibition. Another possibility is that due to the open conformation of the binding site on the C1 domain, Gαi can bind to the AC5+ATP+Gαs complex to form a ternary complex, whose existence is still unknown.

## Conclusions

We perform all-atom molecular dynamics simulations in an attempt to better understand the regulation of adenylyl cyclase, a key enzymatic player in cellular signalling cascades. Microsecond-scale simulations of the G-protein subunit Gαi bound to adenylyl cyclase in the presence and in the absence of ATP in two different conformations help to better understand some features of this important signal transmission protein since no structural information on this complex is available. They notably provide information on a single, non-chimeric adenylyl cyclase isoform, AC5, bound to the inhibitory G-protein in the presence and in the absence of ATP.

The simulations show that protein binding creates significant changes in the structure and in flexibility, throughout AC5 and due to a strong allosteric coupling existing within AC5 in a different fashion than the stimulatory G-protein and ATP. They provide data that help to explain the inhibition action of Gαi, whose binding increases the conformational and positional fluctuations of ATP in the active site of AC5 and its flexibility by moving away the key residues involved in the enzymatic reactions.

Our results also show that Gαi binding to the C1 domain does not impact C1/C2 interface complementarity, flexibilizes C1 domain and significantly closes the angle between the C2 α-helices that cannot bind Gsα when Gαi is in tilted conformation. The simultaneous binding of ATP and Gαi in a titled conformation at the AC5 interface results in a rigidification of the C2 domain, without affecting the C1/C2 interface complementarity, and a slight increase of the angle between the C2 α-helices. Hence, Gαi has an important impact on AC5 dynamics and its effects are enhanced when ATP binds, by increasing the conformational freedom of the bound ligand, thus putting it in an unfavourable configuration within its binding site and not allowing to establish key interactions between ATP and AC5 that leave therefore AC5 completely inactive.

Our simulations also show that ATP has a crucial role in the regulation of AC5 and we cannot exclude the presence of ATP during the inhibition. Our previous simulations already showed that ATP binding could influence the binding of the inhibitory G-protein subunit αi at the domain C1. Here, we propose that the presence of ATP is needed to induce the competition between Gsα and Gαi to tightly regulate AC5. However, based on our results, we cannot exclude the existence of the Gαi+AC5+ATP+Gsα complex and the Gαi+AC5+Gsα complex. For the latter, other molecular dynamic studies of this hypothetical ternary complex concluded that if existing it would be inactive [29,30]. Further studies have to be conducted to shed lights on this point.

## Methods

### Models

Models of the cytoplasmic domains of AC5 and Gαi protein were built by homology to known proteins using Modeller v9.12 [42]. In each case, 100 homology models were generated and the model with the lowest DOPE score [43] was selected. For AC5, a homology model for the mouse sequence with bound ATP was generated using with structure 1CJK [13] as template. The identity percentage between template and model is equal to 98% for the C1 domain and 57% for the C2 domain. Since we observed in our earlier study [25] that the conformation of AC5 is affected by the binding of G proteins, we used the structure closest to the centre of the largest cluster of the last 500 ns of the MD simulation of AC5+ATP [25], in the absence of G proteins.

For Gαi, a homology model for the mouse sequence bound with GSP and Mg ion was generated using Modeller with three templates: 1CJK [13] (bovine Gsα 38% sequence identity), 1AS3 [44] (rat Gαi, 81% identity), and 1AGR [45] (rat Gαi, 87% sequence identity). This model was used in docking (see below). In addition, we considered a model of Gαi sampled from MD simulation: a simulation of 1 µs was run starting from the homology model. The structure closest to the center of the largest cluster observed during the simulation was used for docking.

Models of AC5 and Gαi were docked using CLUSPRO [46–48] to generate Gαi+AC5+ATP complexes shown in Figure 2, using known interface residues on both proteins as restraints (see Figure S12). A first complex was built by docking the model of AC5 sampled from simulation with the homology model of Gαi. The resulting complex locates Gαi in an orientation similar to Gsα with respect to AC5 in the 1CJK complex, with the G protein binding in the groove formed by the two α-helices (see Figure 2A). This orientation is called the symmetrical orientation. Resulting complex is denoted Gαi_sym+AC5+ATP.

A second complex was built by docking the model of AC5 sampled from MD simulation with the model of Gαi sampled from MD simulation. In the resulting complex, Gαi is tilted compared to the Gsα orientation with respect to AC5: the G protein is in contact not only with the helix groove, but also with residues on the side of the C1 domain (see Figure 2B). Resulting complex is denoted Gαi_tilted+AC5+ATP.

Available structures of Gαi display different conformations of the N-terminal helix: either protruding from the structure (in 1AGR) or packed onto the structure core (in 1AS3). In the initial model, this helix was packed. During the simulation of Gαi, this helix appeared very mobile. In order to minimize possible bias and to avoid Periodic Boundary Condition problems in the simulations of Gαi+AC5 complexes, we manually unpacked the N-terminal helix from the structure core after the docking step.

Systems without ATP were also simulated, starting from the same systems after ATP removal. Throughout the study, we compared our results with those obtained in our previous study of AC5 alone and in complex with the activating protein Gsα [25] (see Figure 1, right box).

### All-atom molecular dynamics simulations

Molecular dynamics simulations were performed with the GROMACS 5 package [33–36,49] using the Amber99SB-ILDN force field for proteins that has been shown to yield an accurate description of many structural and dynamical properties of proteins [38,50–52]. Side chain protonation states of titratable amino acids were assigned using a value of pH = 7.4 with the help of the pdb2pqr software [53]. Capping acetyl and methyl-amino groups were added to the N and C termini of both AC5 domains and Gαi. The four states we study (Gαi_sym+AC5, Gαi_sym+AC5+ATP, Gαi_tilted+AC5, Gαi_tilted+AC5+ATP) were each placed in a truncated octahedral box and solvated with TIP3P water molecules [54] to a depth of at least 11 Å. The solute was neutralized with potassium cations and then K+Cl-ion pairs [55] were added to reach a physiological salt concentration of 0.15 M. Parameters for ATP and GTP were taken from [56]. The parameters for Mg^2+^ came from [57]. This new set of parameters was developed to improve the kinetic properties of Mg^2+^ ions with water and with the phosphate ion and it was implemented in Amber99. This new set of parameters also provided a better description of the structure of Mg^2+^-phosphate binding than previous sets (these interactions are naturally important in our simulations in the presence of ATP) [57]. Hence, the combination of Amber 99SB-ILDN and the new set of parameters of Mg^2+^ ions is currently the best choice to reproduce the dynamics of AC5 and Gsα, and to properly describe the interactions of Mg^2+^ with AC5 and ATP.

Long-range electrostatic interactions were treated using the particle mesh Ewald method [58,59] with a real-space cutoff of 10 Å. We used virtual interaction sites for the hydrogens and bond lengths were restrained using P-LINCS [36,60], allowing a time step of 4 fs [61]. Translational movement of the solute was removed every 1000 steps to avoid any kinetic energy build-up [62]. After energy minimization of the solvent and equilibration of the solvated system for 10 ns using a Berendsen thermostat (t_T_ = 1 ps) and Berendsen pressure coupling (t_P_ = 4 ps) [63], the simulations were carried out in an NTP ensemble at a temperature of 310 K and a pressure of 1 bar using a Bussi velocity-rescaling thermostat [64] (t_T_ = 1 ps) and a Parrinello-Rahman barostat (t_P_ = 1 ps) [65]. Simulations were carried out using typically between 72 and 120 computer cores depending on the system size, which allowed a production rate of about 100 ns/ day. Analysis was carried out on a 1.1 µs production segment for each simulation, following a 400 ns equilibration period as in our previous study [25].

### Analysis of all-atom MD simulations

We analyzed our all-atom MD simulations using average structures, time-averaged properties such as RMSD (Root-Mean-Square-Deviation), angle between helices, distance between helix axes, distance between the ATP/Mg^2+^ ion and some key residues, and specific geometrical measurements described below, protein-protein and protein-ligand interface characteristics and, in some cases, residue-by-residue conformational and dynamic properties.

When RMSD distributions indicated the existence of distinct conformations, a cluster analysis was carried out using the gromos algorithm of GROMACS [66], using a RMSD cutoff equal to 1.5 Å on backbone atoms, on the conformations collected in the production phase. Clusters accounting for less than 100 ns were discarded.

The C1/C2 interface was characterized using three quantities: the gap volume, the change of accessible surface area upon binding (ΔASA), and the Gap index [67,68]. The Gap index, defined by the gap volume between two protein chains divided by the interface area, measures the shape complementarity at protein-protein interfaces [68]. The gap volume was computed by the SURFNET software [67], and the interface area was calculated using a local implementation of the Lee and Richards algorithm [69] and the same radii.

In order to characterize G protein binding sites, as in our previous work [25], we computed the angle α_C2_ between the pairs of α-helices within domain C2 that bind Gsα (termed α3 and α4 in Figure 3) and also the angle α_C1_ between the quasi-symmetric pair of helices within domain C1 (termed α1 and α2 in Figure 3) that binds Gαi in the present study. The angles were measured using helical axes derived from the residues that remain in stable α-helical conformations throughout the simulations (C1: 408–420 and 468–475, C2: 910–918 and 978– 988) as defined in [25]. We also computed the distances between the center of the helices in each domain (d_C1_ and d_C2_, respectively).

To characterize protein-protein interfaces, we computed the interface contacts with the python/C code available at https://github.com/MMSB-MOBI/ccmap, using a fixed cutoff of 5 Å between heavy atoms.

To characterize the ATP binding site, we computed two distances between ATP and two key residues for AC5 activity (distance between O_2_γ and Lys 1065 and between O_2_α and Arg 1029) and the distance between Asp 460 and Asp396 and the two Mg^2+^ ions.

## Supporting Information

Supporting Information including 3 tables and 12 figures.

## Acknowledgements

This project/research has received funding from the European Union’s Horizon 2020 Framework Programme for Research and Innovation under the Specific Grant Agreement No. 720270 (Human Brain Project SGA1) and the Specific Grant Agreement No. 785907 (Human Brain Project SGA2). We thank Alexis Michon and Samuel Bosquin (UMS 3760, Institut de Biologie et Chimie des Protéines, Lyon, France) for technical assistance and hardware support. We wish to acknowledge GENCI for a generous allocation of computer time on the CINES supercomputer OCCIGEN (Grant 2016 - c201607758, Grant 2017 - A0020707585, Grant 2018 - A0040710357).

## Author Contributions

**Conceptualization:** Elisa Frezza, Juliette Martin.

**Data curation:** Elisa Frezza, Juliette Martin

**Formal analysis:** Tina Méryl-Amans, Juliette Martin.

**Investigation:** Elisa Frezza, Tina Méryl-Amans

**Methodology:** Elisa Frezza, Juliette Martin.

**Validation:** Elisa Frezza, Juliette Martin.

**Supervision:** Elisa Frezza, Juliette Martin.

**Writing – original draft:** Elisa Frezza, Juliette Martin.

**Writing – review & editing:** Elisa Frezza, Juliette Martin.

## SUPPLEMENTARY MATERIAL

### S1. Figures

**Figure S1.**
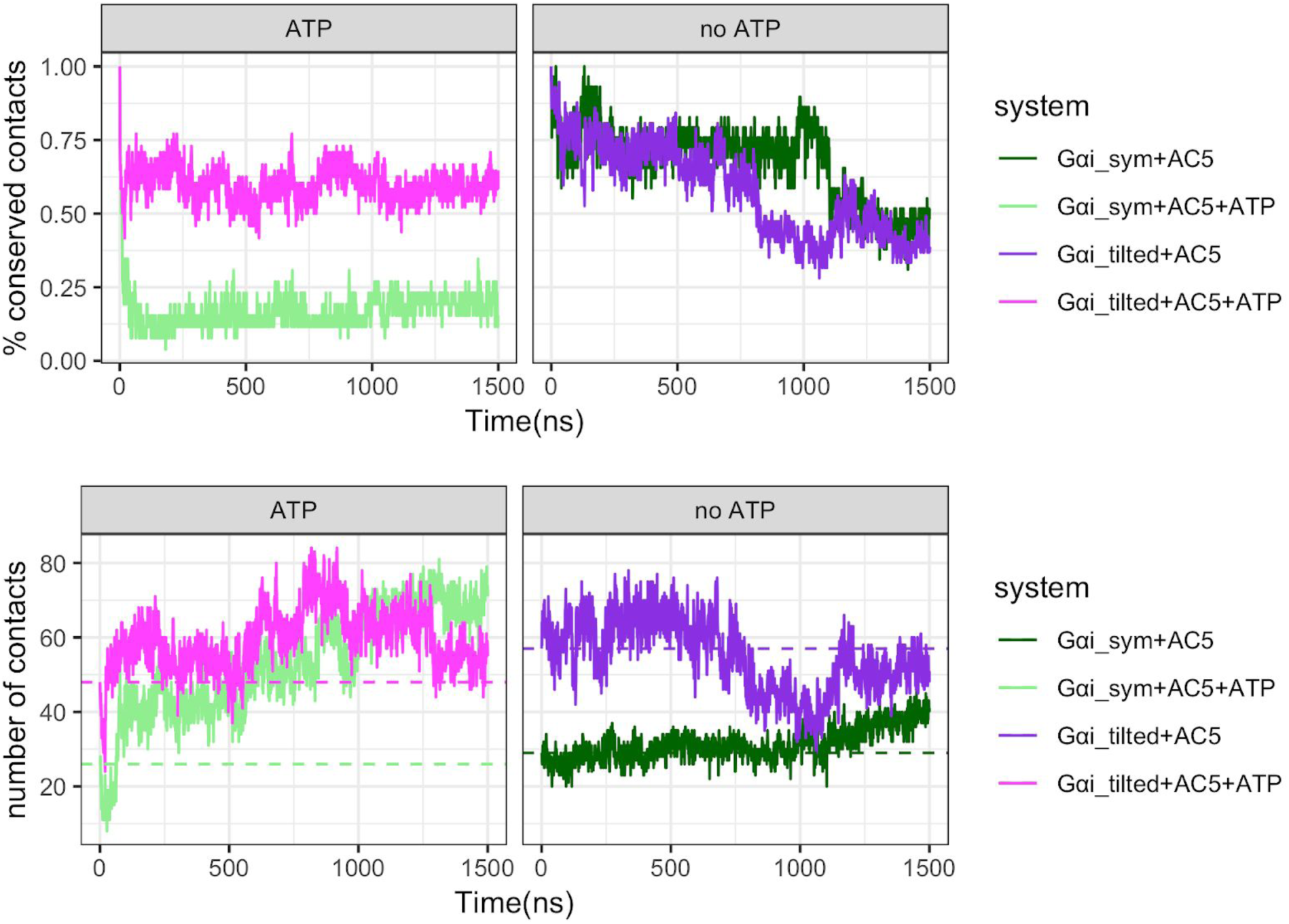
Number of contacts at the AC5/Gαi interface. Contacts are defined using a 5 Å cut-off between heavy atoms. Top row: fraction of initial contacts (T=0) that are maintained as a function of time. Bottom row: total number of contacts between AC5 and Gαi, dashed horizontal lines indicate the number of contacts at T=0.

**Figure S2.**
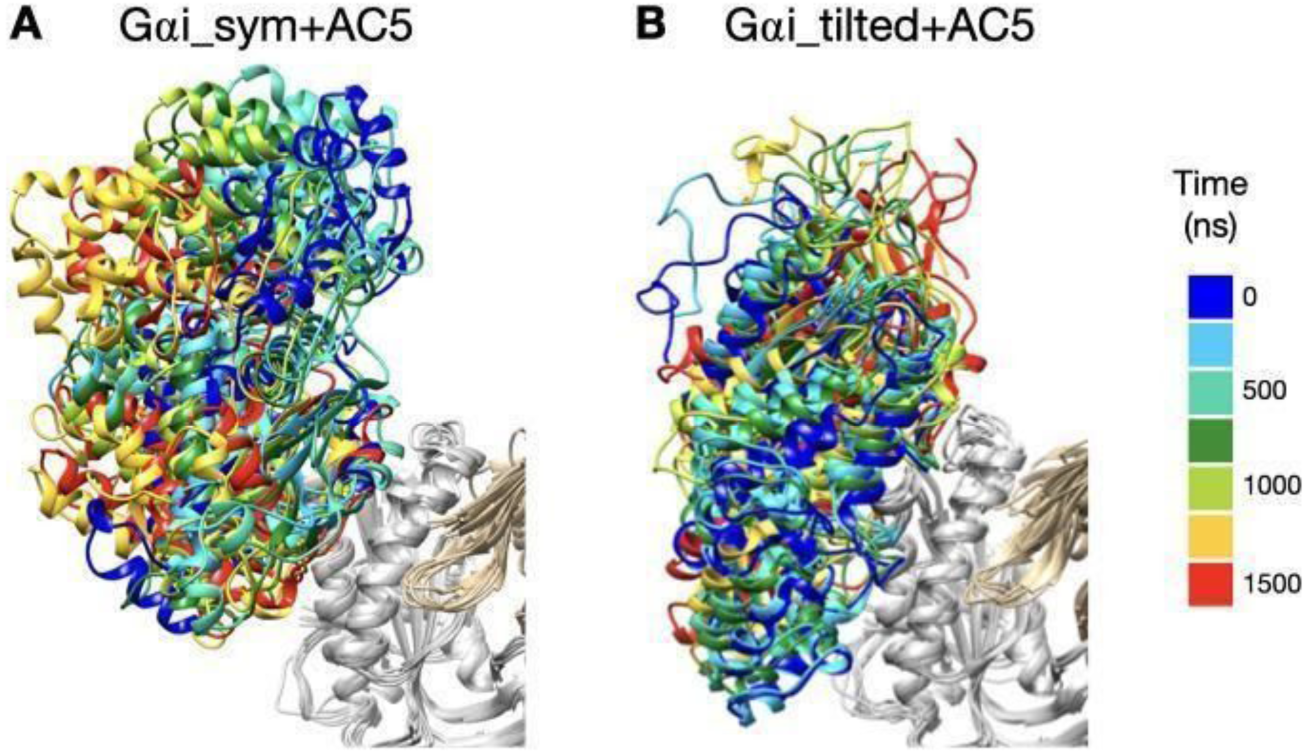
Snapshots of the Gαi+AC5 complexes observed during the simulations without ATP, viewed from the membrane side. Structures extracted every 250 ns are colored on a rainbow scale from blue to red. The C1 domain of AC5 is colored in grey and the C2 domain in beige.

**Figure S3:**
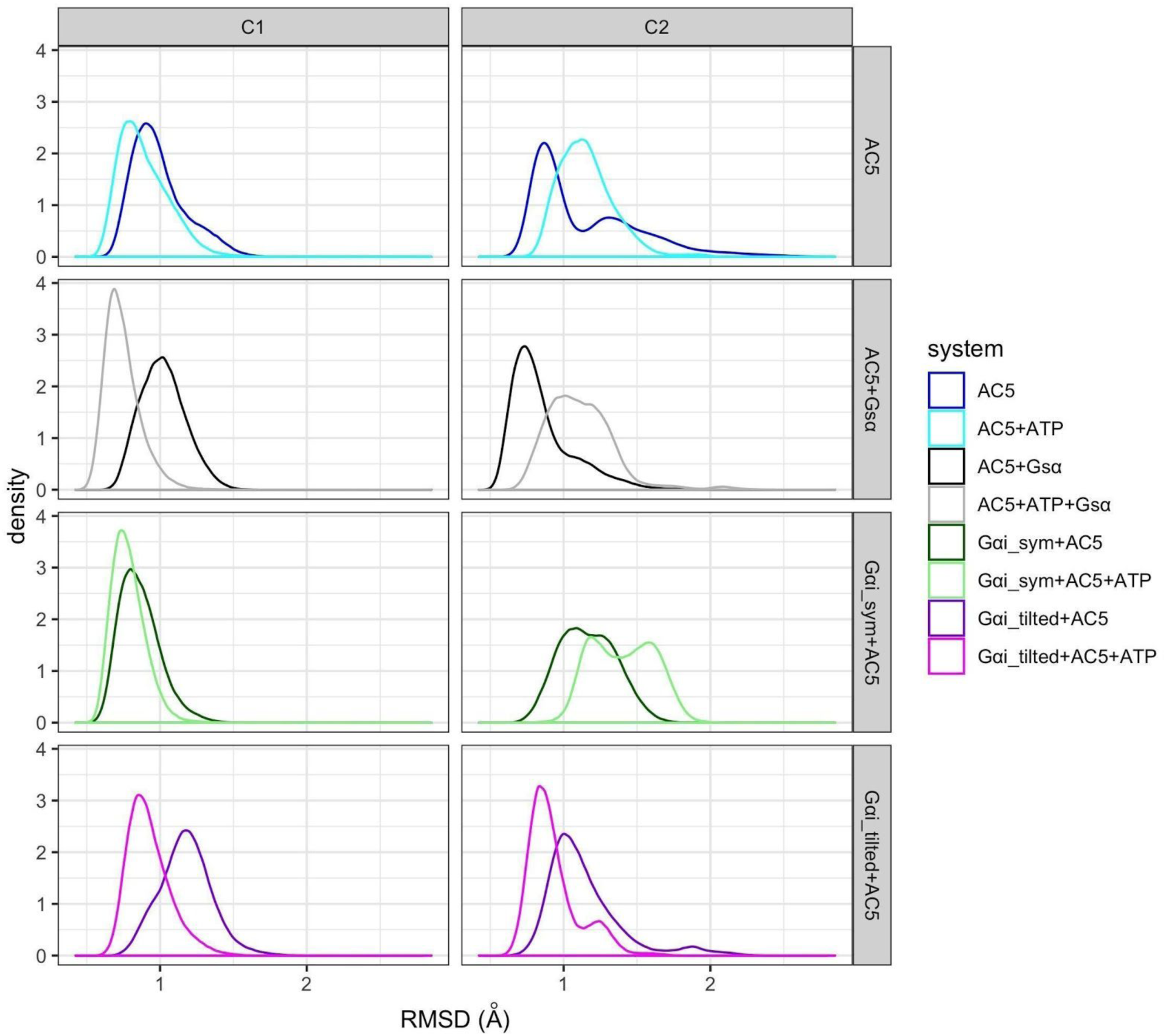
RMSD distribution for C1 and C2 domains, with respect to average structures, in the simulations with and without ATP. Data for AC5 and AC5+Gsα, with and without ATP, taken from (Frezza E, Martin J, Lavery R. A molecular dynamics study of adenylyl cyclase: The impact of ATP and G-protein binding. PLOS ONE. 2018;13: e0196207. doi:10.1371/journal.pone.0196207;Frezza E, Martin J, Lavery R. A molecular dynamics study of adenylyl cyclase: the impact of ATP and G-protein binding. Zenodo; 2018. doi:10.5281/zenodo.1213125).

**Figure S4:**
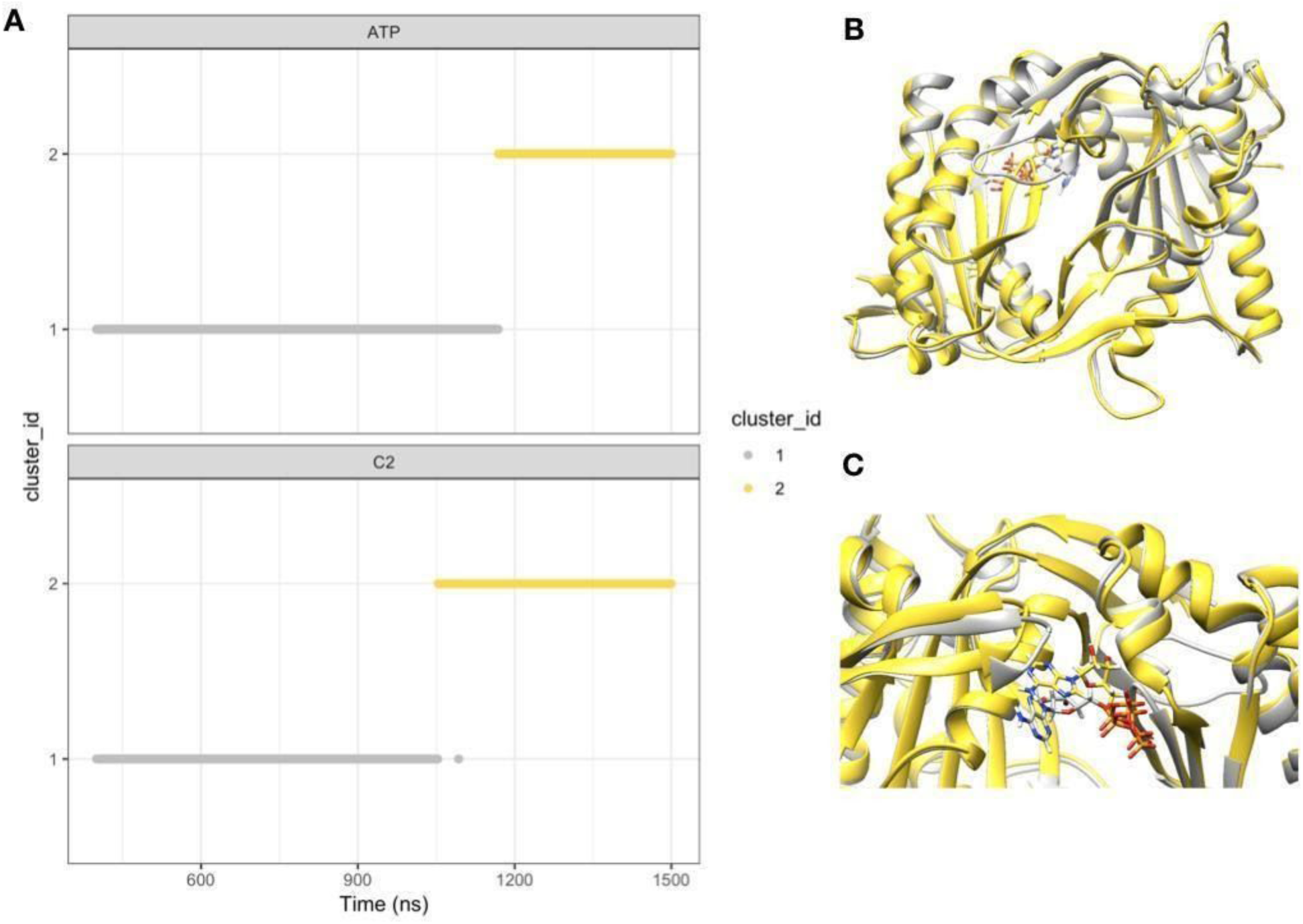
Substates of ATP and C2 observed in the Gαi_sym+AC5+ATP simulation. A: structural clusters obtained using gromos (cutoff=1.5 Å on backbone atoms for C2 and cutoff=1 Å for ATP), B: average structures viewed from the membrane side, C: close-up view on the β4 loop, from the cytoplasmic side. The average structures are colored in grey (400ns<T<1100 ns) and yellow (1200ns<T<1500ns).

**Figure S5:**
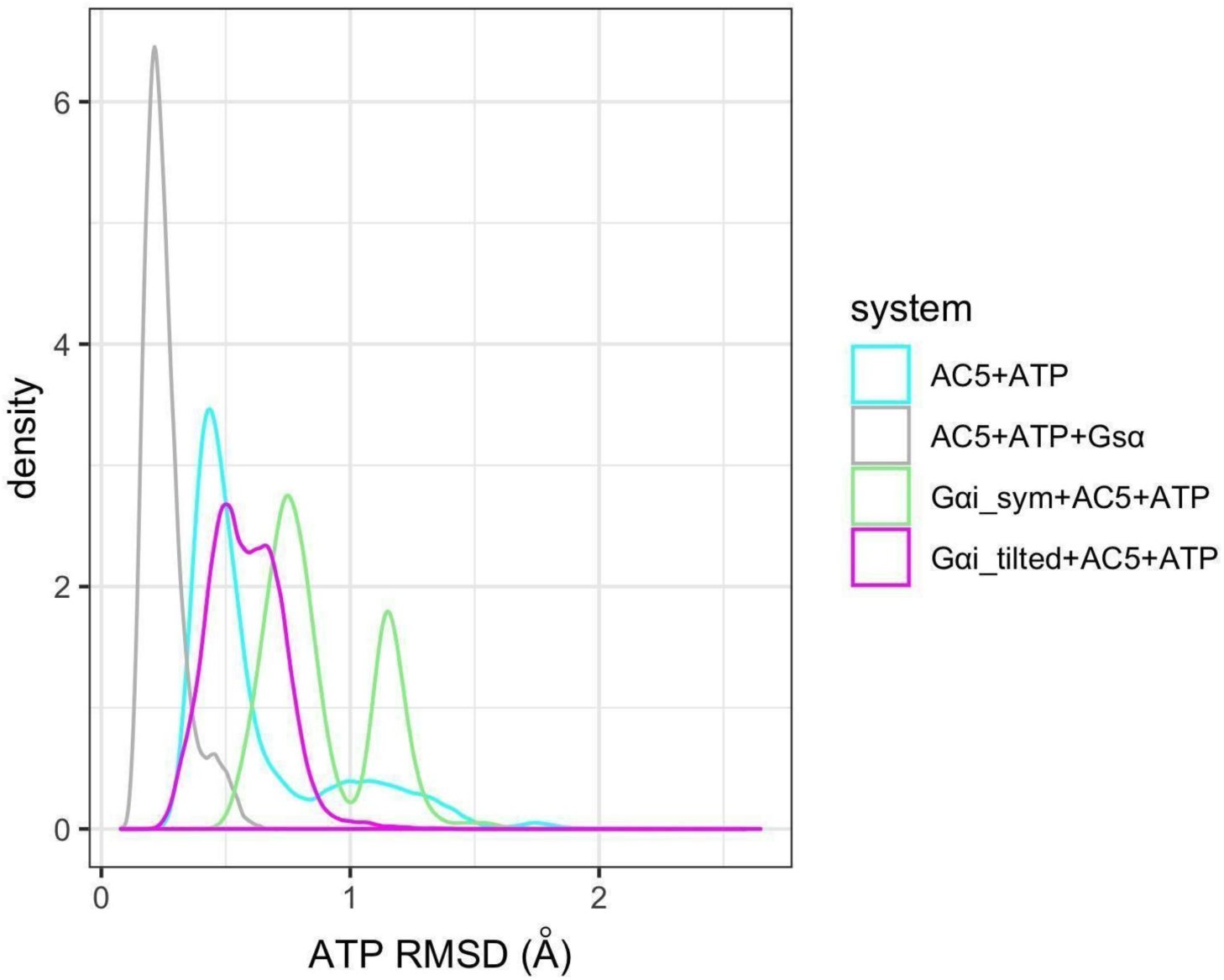
RMSD distribution for ATP. Data for AC5+ATP and AC5+ATP+Gsα taken from (Frezza E, Martin J, Lavery R. A molecular dynamics study of adenylyl cyclase: The impact of ATP and G-protein binding. PLOS ONE. 2018;13: e0196207. doi:10.1371/journal.pone.0196207;Frezza E, Martin J, Lavery R. A molecular dynamics study of adenylyl cyclase: the impact of ATP and G-protein binding. Zenodo; 2018. doi:10.5281/zenodo.1213125).

**Figure S6:**
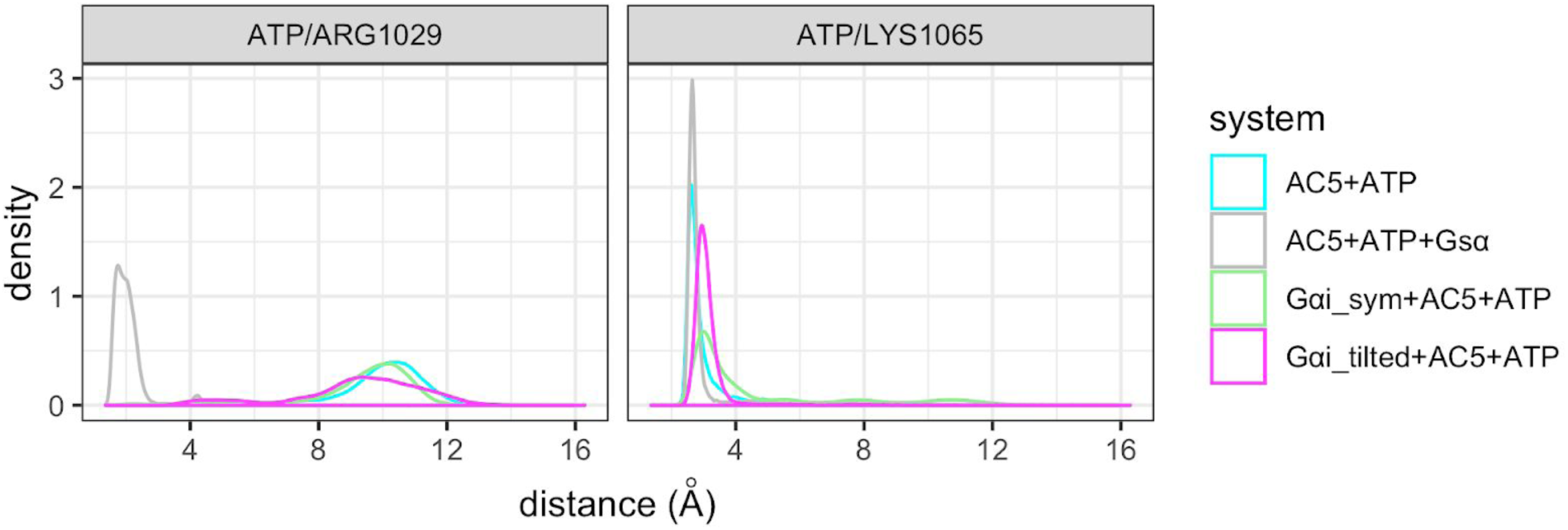
Distance between ATP and key residues of C2. Data for AC5+ATP and AC5+ATP+Gsα are from our previous study (Frezza E, Martin J, Lavery R. A molecular dynamics study of adenylyl cyclase: The impact of ATP and G-protein binding. PLOS ONE. 2018;13: e0196207. doi:10.1371/journal.pone.0196207;Frezza E, Martin J, Lavery R. A molecular dynamics study of adenylyl cyclase: the impact of ATP and G-protein binding. Zenodo; 2018. doi:10.5281/zenodo.1213125). ATP/ARG1029: distance between the O2α of ATP and the center of mass of terminal hydrogen atoms which are covalently bound to Nε of ARG1029. ATP/LYS1065: distance between the O2γ of ATP and the center of mass of the terminal hydrogen atoms which are covalently bound to Nζ of Lys 1065.

**Figure S7:**
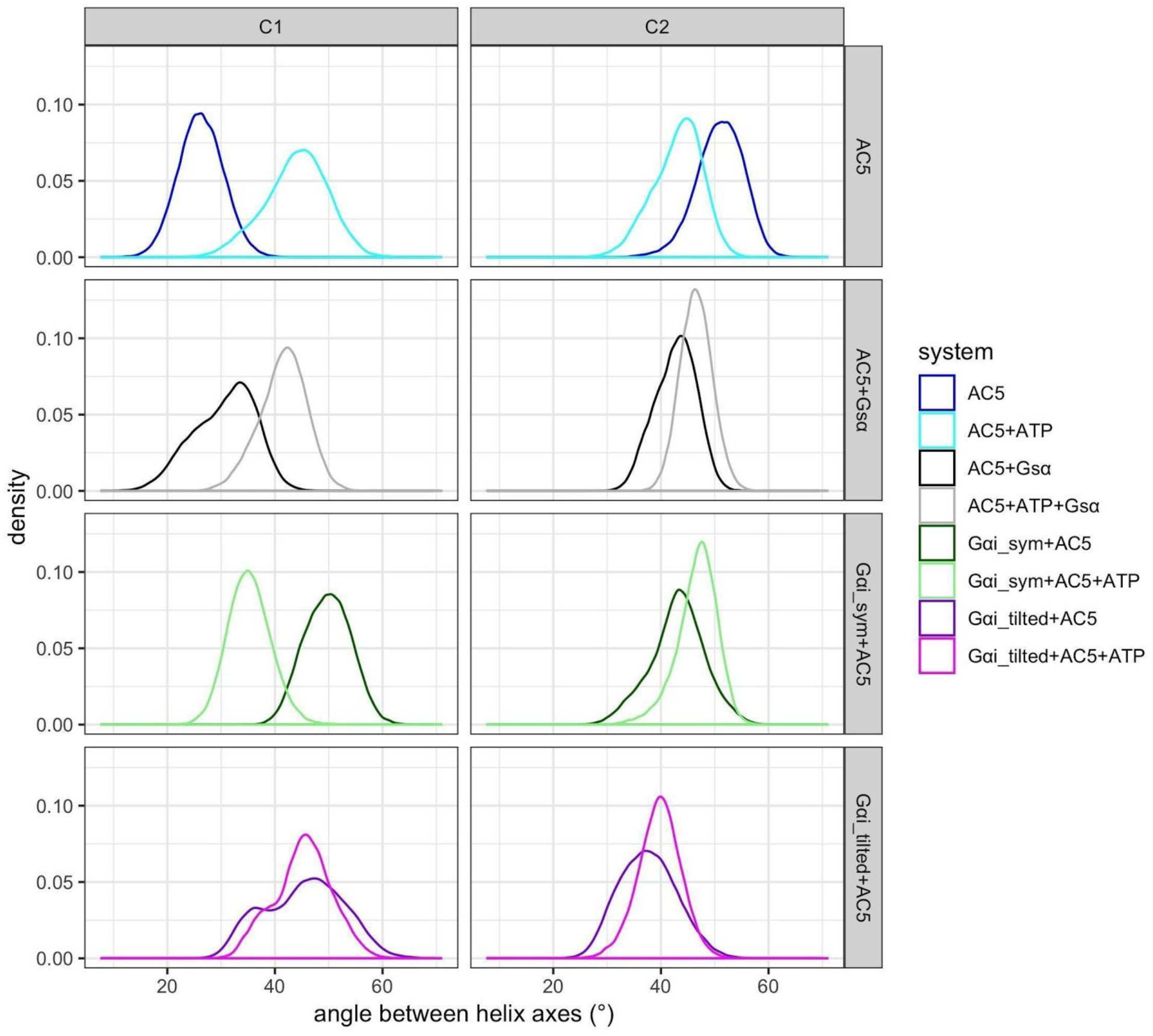
Distribution of the angles between helix axes. Data for AC5 and AC5+Gsα, with and without ATP, taken from (Frezza E, Martin J, Lavery R. A molecular dynamics study of adenylyl cyclase: The impact of ATP and G-protein binding. PLOS ONE. 2018;13: e0196207. doi:10.1371/journal.pone.0196207;Frezza E, Martin J, Lavery R. A molecular dynamics study of adenylyl cyclase: the impact of ATP and G-protein binding. Zenodo; 2018. doi:10.5281/zenodo.1213125).

**Figure S8:**
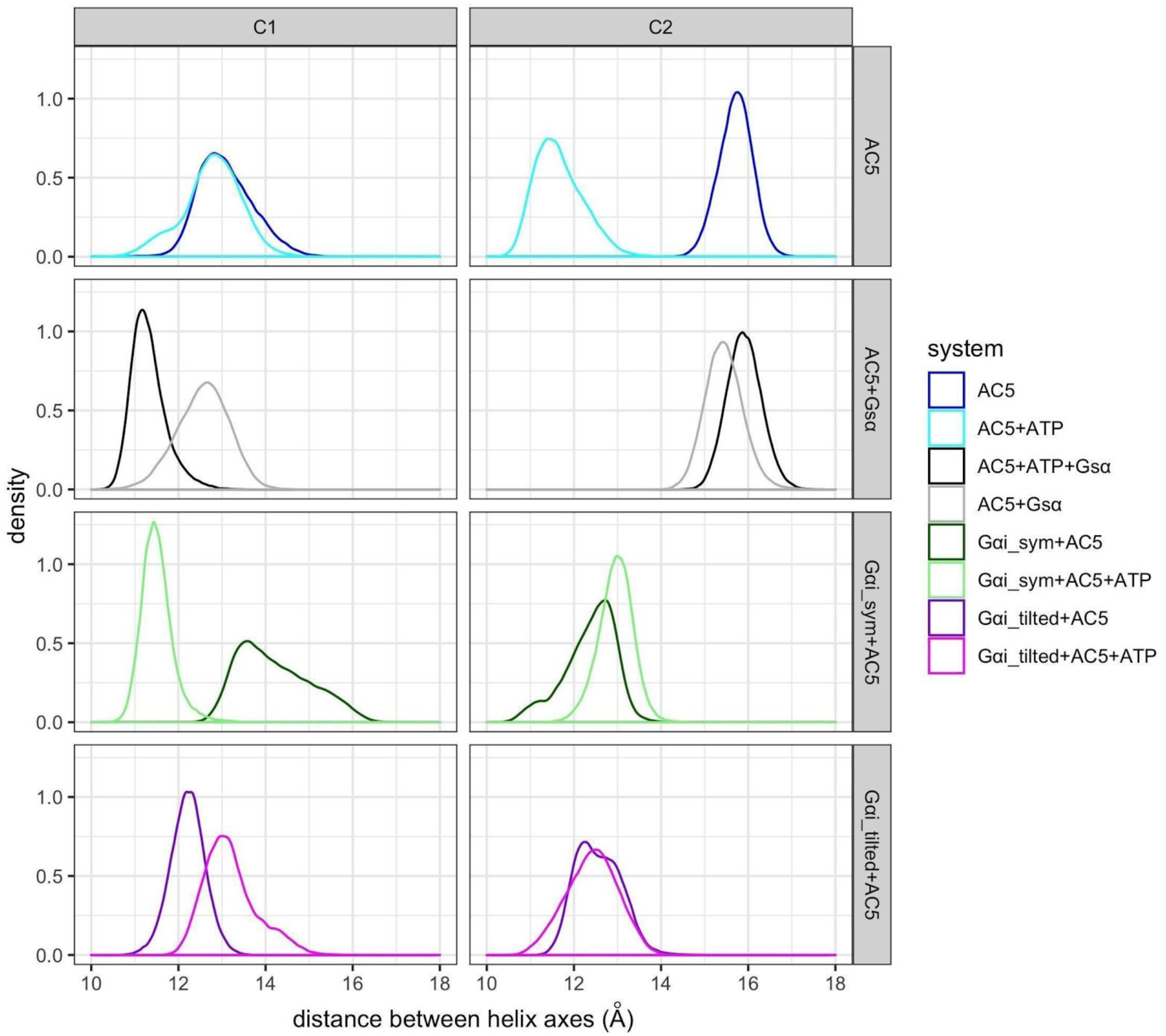
Distribution of the distance between helix axes. Data for AC5 and AC5+Gsα, with and without ATP, taken from (Frezza E, Martin J, Lavery R. A molecular dynamics study of adenylyl cyclase: The impact of ATP and G-protein binding. PLOS ONE. 2018;13: e0196207. doi:10.1371/journal.pone.0196207;Frezza E, Martin J, Lavery R. A molecular dynamics study of adenylyl cyclase: the impact of ATP and G-protein binding. Zenodo; 2018. doi:10.5281/zenodo.1213125).

**Figure S9:**
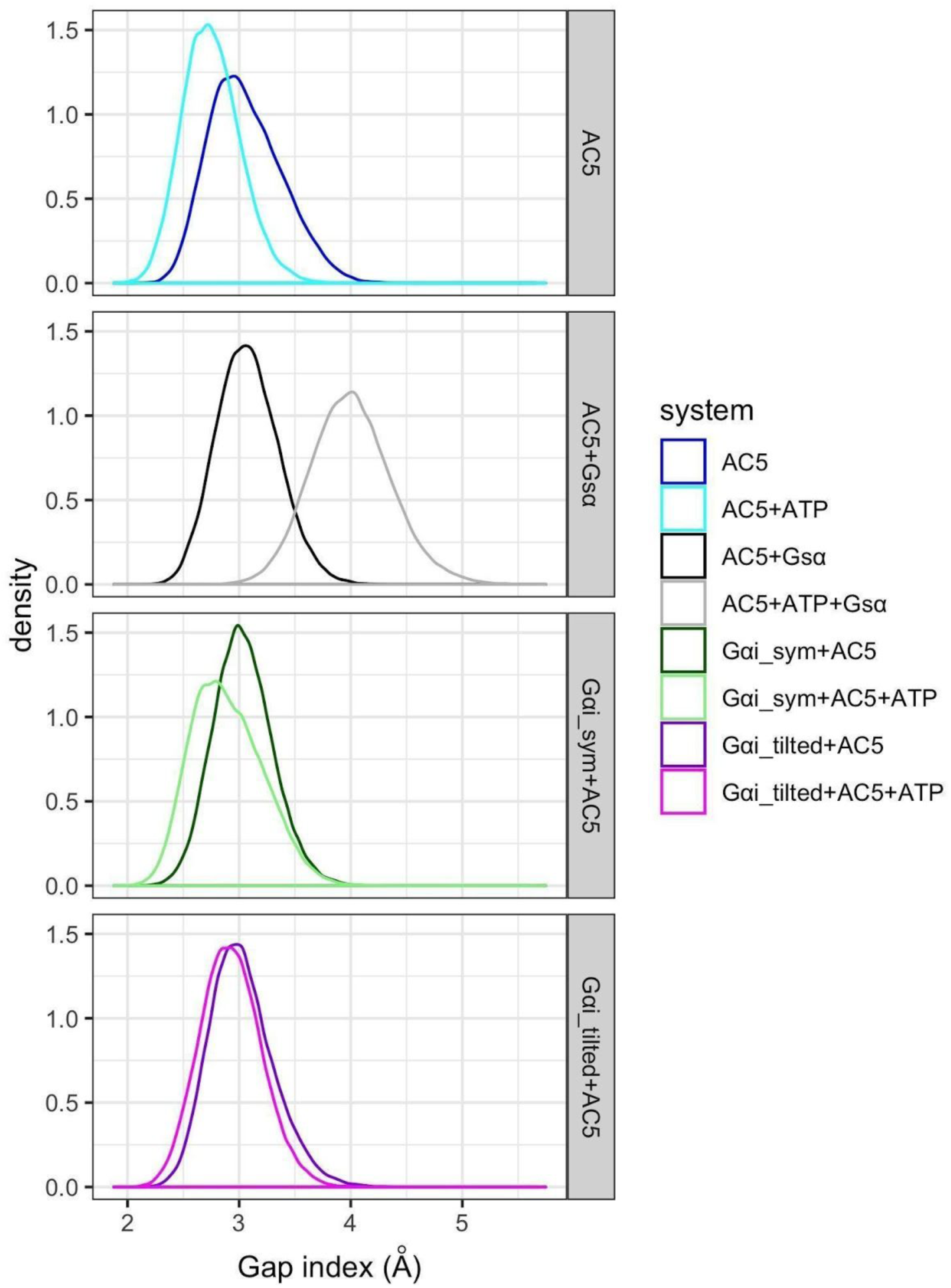
Distribution of the C1/C2 interface Gap index. Data for AC5 and AC5+Gsα, with and without ATP, taken from (Frezza E, Martin J, Lavery R. A molecular dynamics study of adenylyl cyclase: The impact of ATP and G-protein binding. PLOS ONE. 2018;13: e0196207. doi:10.1371/journal.pone.0196207;Frezza E, Martin J, Lavery R. A molecular dynamics study of adenylyl cyclase: the impact of ATP and G-protein binding. Zenodo; 2018. doi:10.5281/zenodo.1213125).

**Figure S10.**
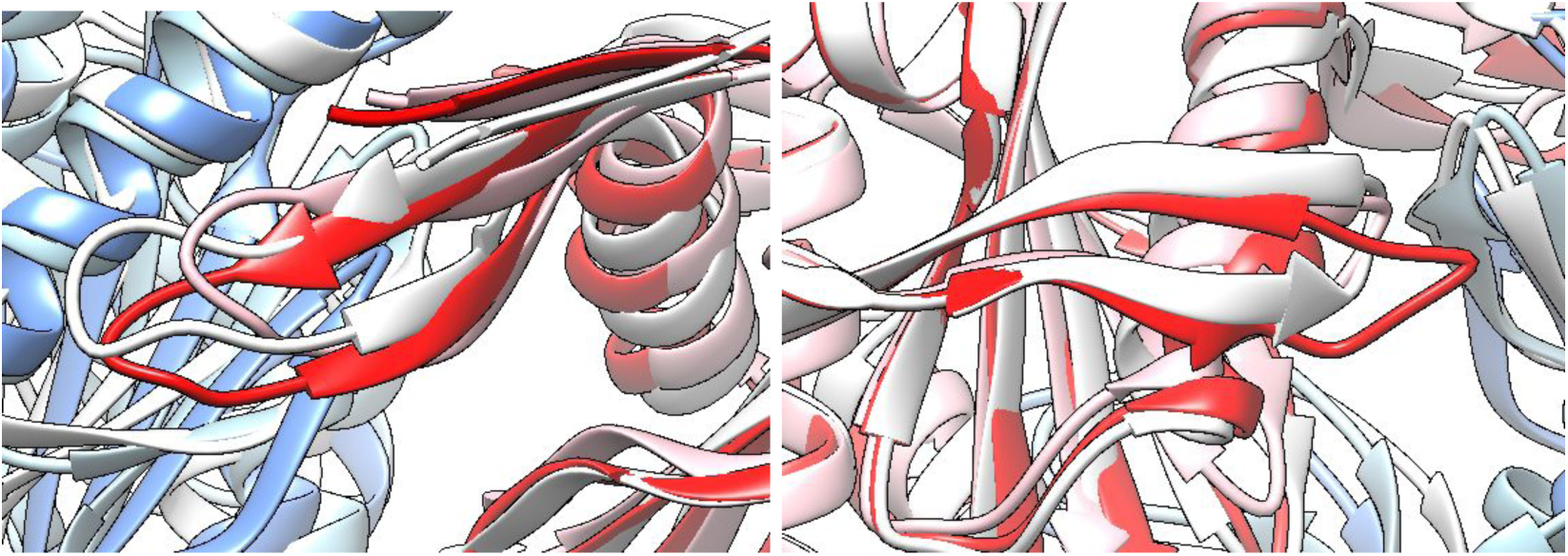
Local comparison of C2 loops in average structures without ATP. White: AC5, red/blue: Gαi_tilted+AC5, pink/cyan: Gαi_sym+AC5. Left panel: β2 loop, right panel: β4 loop.

**Figure S11.**
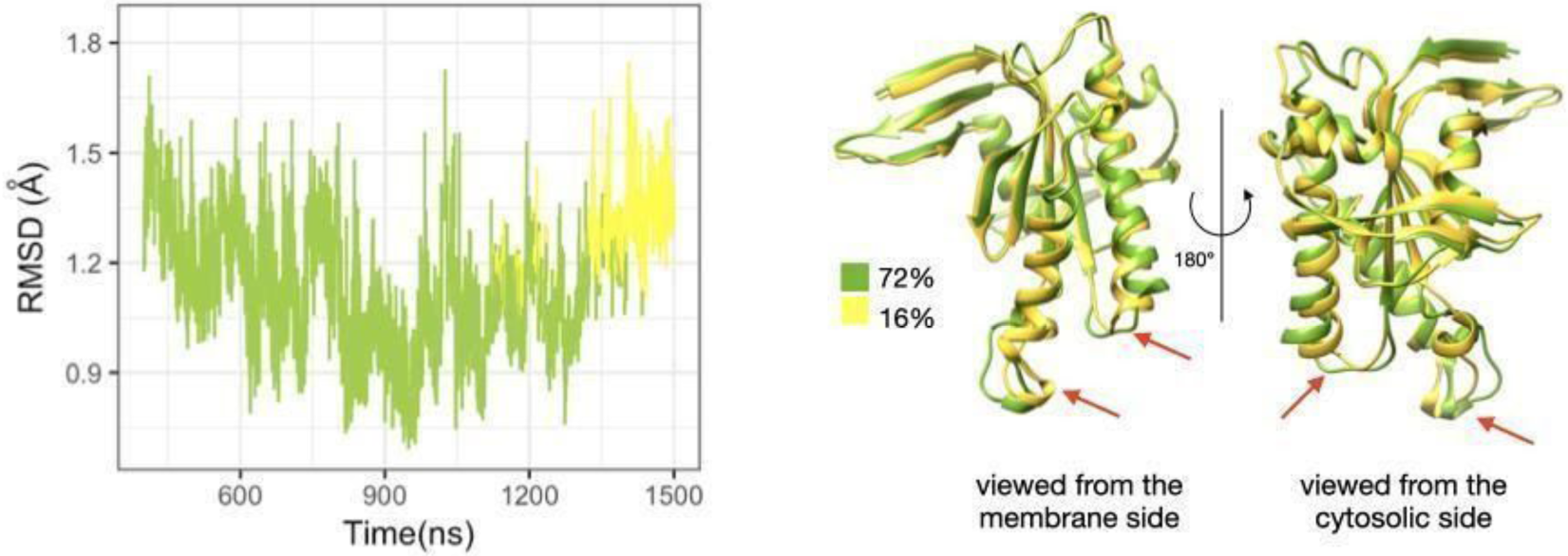
Sub-states of domain C2 observed during the simulation of Gαi_sym+AC5 complex, without ATP. Left: RMSD time series for the C2 domain, colored according to cluster membership. Right: structures closest to the center of each cluster, and relative size of each cluster as percentages. Prominent structural changes are indicated by red arrows.

**Figure S12:**
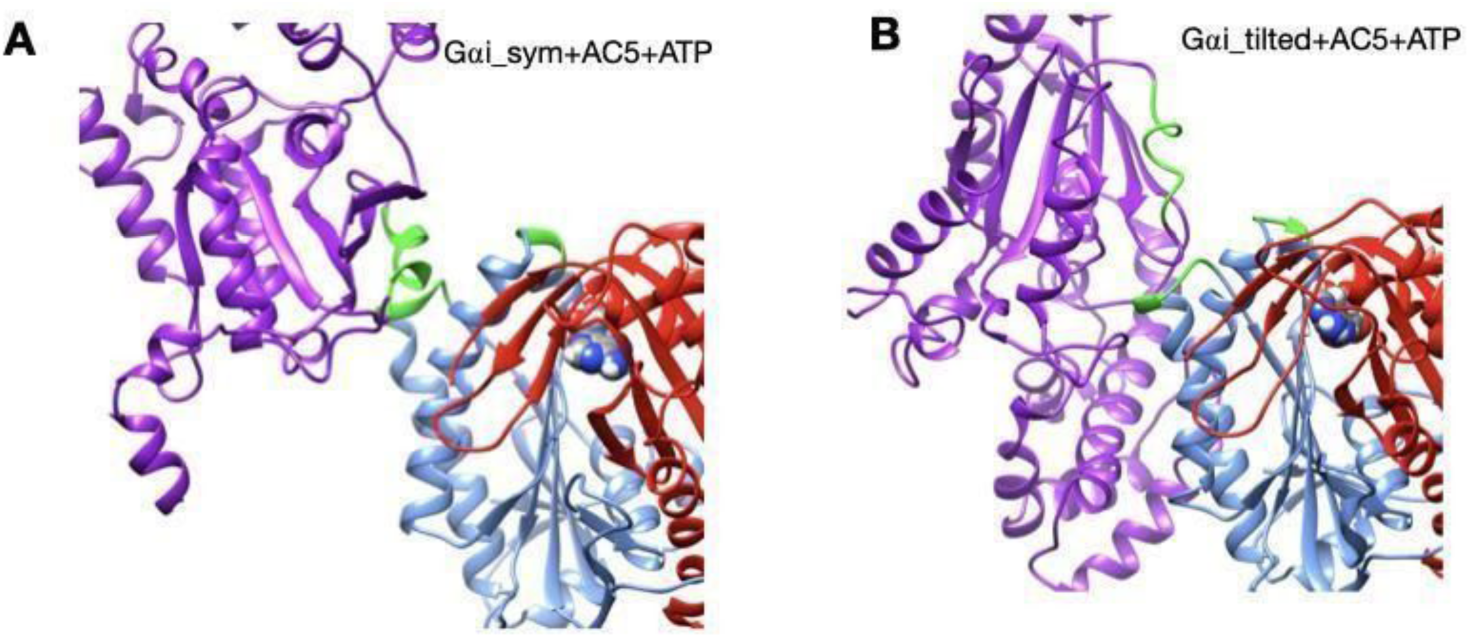
Gαi+AC5 complexes, viewed from the membrane side. Gαi is colored in purple, AC5 in blue (C1 domain) and red (C2 domain) and interface residues used as restraints for docking are colored in green: residues 101-105 and 31-33 in AC5, and residues 202-209 in Gαi.

### S2. Tables

**Table S1.** Average and standard deviation of backbone RMSD for the C1 and C2 domains of AC5, angle between helices (α_C1_ and α_C2_), distance between helices axes (d_C1_ and d_C2_), Gap index for the interface C1/C2. ^a^: data in italic are from our previous study (Frezza E, Martin J, Lavery R. A molecular dynamics study of adenylyl cyclase: The impact of ATP and G-protein binding. PLOS ONE. 2018;13: e0196207. doi:10.1371/journal.pone.0196207;Frezza E, Martin J, Lavery R. A molecular dynamics study of adenylyl cyclase: the impact of ATP and G-protein binding. Zenodo; 2018. doi:10.5281/zenodo.1213125).

| System | RMSD C1 (Å) | RMSD C2 (Å) | $\alpha_{C1}$ (°) | $d_{C1}$ (Å) | $\alpha_{C2}$ (°) | $d_{C2}$ (Å) | Gap index C1/C2 (Å) | RMSD ATP (Å) |
| --- | --- | --- | --- | --- | --- | --- | --- | --- |
| AC5+ATP<sup>a</sup> | *0.9 ± 0.2* | *1.2 ± 0.2* | *44 ± 6* | *12.8 ± 0.7* | *43 ± 5* | *11.7 ± 0.6* | *2.8 ± 0.3* | *0.6 ± 0.3* |
| AC5+ATP+Gsa<sup>a</sup> | *0.8 ± 0.1* | *1.1 ± 0.2* | *42 ± 5* | *11.3 ± 0.4* | *47 ± 3* | *15.9 ± 0.4* | *3.8 ± 0.5* | *0.3 ± 0.1* |
| Gai_sym+AC5+ATP | 0.8 ± 0.1 | 1.4 ± 0.2 | 35 ± 4 | 11.5 ± 0.5 | 47 ± 4 | 13.0 ± 0.4 | 2.9 ± 0.3 | 0.9 ± 0.2 |
| Gai_tilted+AC5+ATP | 0.9 ± 0.1 | 0.9 ± 0.2 | 45 ± 5 | 13.2 ± 0.6 | 40 ± 4 | 12.4 ± 0.6 | 2.9 ± 0.3 | 0.6 ± 0.1 |
| AC5<sup>a</sup> | *1.0 ± 0.2* | *1.2 ± 0.4* | *26 ± 4* | *13.1 ± 0.6* | *50 ± 4* | *15.7 ± 0.4* | *3.1 ± 0.3* |  |
| AC5+Gsa<sup>a</sup> | *1.0 ± 0.2* | *0.9 ± 0.2* | *31 ± 6* | *12.6 ± 0.6* | *43 ± 4* | *15.4 ± 0.4* | *3.1 ± 0.3* |  |
| Gai_sym+AC5 | 0.9 ± 0.1 | 1.2 ± 0.2 | 50 ± 4 | 14.2 ± 0.8 | 43 ± 5 | 12.4 ± 0.6 | 3.0 ± 0.3 |  |
| Gai_tilted+AC5 | 1.2 ± 0.2 | 1.1 ± 0.3 | 45 ± 7 | 12.2 ± 0.4 | 38 ± 5 | 12.6 ± 0.5 | 3.0 ± 0.3 |  |

**Table S2:** Mean values and standard deviation of Gap index for the Gαi/AC5 interface. ^a^: data in italic are from our previous study for the Gap index for the Gsα/AC5 interface.

| System | Gap Volume (Å<sup>3</sup>) | ΔASA (Å<sup>2</sup>) | Gap index (Å) |
| --- | --- | --- | --- |
| Gai_sym+AC5+ATP | 7569 ± 504 | 1413 ± 157 | 5.4 ± 0.7 |
| Gai_tilted+AC5+ATP | 5640 ± 469 | 1345 ± 152 | 4.2 ± 0.6 |
| Gai_sym+AC5 | 2850 ± 449 | 881 ± 101 | 3.2 ± 0.4 |
| Gai_tilted+AC5 | 5561 ± 562 | 1225 ± 138 | 4.6 ± 0.7 |
| AC5+ATP+Gsa<sup>a</sup> | 3286 ± 333 | 1244 ± 123 | 2.7 ± 0.5 |
| *AC5+Gsa<sup>a</sup>* | *3424 ± 361* | *1067 ± 63* | *3.2 ± 0.4* |

**Table S3.**
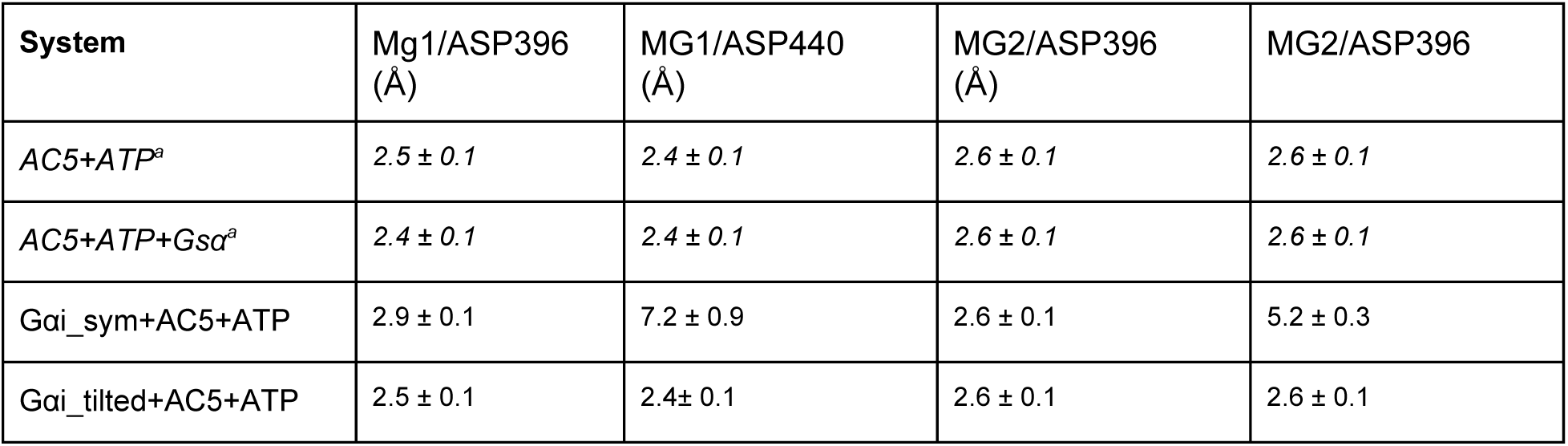
Distance between Mg ions and the arginine residues (Asp 396 and Asp 460 in C1 domain).

